# Gfa, a new integron-borne fosfomycin resistance protein family

**DOI:** 10.64898/2026.10.07.757220

**Authors:** Nicolas Kieffer, Amalia Prieto, Laura Ortiz-Miravalles, Alberto Hipólito, Ester Vergara, Juan Borrero, André Carvalho, José-Antonio Escudero

**Affiliations:** Molecular basis of Adaptation Laboratory, Departamento de Sanidad Animal, Facultad de Veterinaria, Universidad Complutense de Madrid, Madrid, Spain; VISAVET Health Surveillance Centre, Universidad Complutense de Madrid, Spain; Maimonides Biomedical Research Institute of Cordoba, Reina Sofía University Hospital, University of Cordoba, Cordoba, Spain; CIBER de Enfermedades Infecciosas (CIBERINFEC), Instituto de Salud Carlos III (ISCIII), Madrid, Spain; Molecular Basis of Adaptation Laboratory, Departamento de Biotecnología Microbiana, Centro Nacional de Biotecnología, CSIC, Madrid, Spain; Departamento de Biología Molecular, Facultad de Biología, Universidad Autónoma de Madrid, Madrid, Spain; Sección Departamental de Nutrición y Ciencia de los Alimentos (SD-NUTRYCIAL), Facultad de Veterinaria, Universidad Complutense de Madrid, Madrid, Spain

## Abstract

Fosfomycin is increasingly used to treat multidrug-resistant Gram-negative infections, yet resistance surveillance largely relies on homology-based detection of known *fos* genes. Here, we identify Gfa, a previously unrecognized integron-borne fosfomycin resistance determinant uncovered through functional screening of 129 integron gene cassettes. Gfa confers high-level fosfomycin resistance across Enterobacterales and *Pseudomonas aeruginosa* and is resistant to phosphonoformate inhibition. Structural modelling and mutagenesis reveal a conserved cysteine-rich, zinc-organized architecture essential for activity and distinct from canonical Fos enzymes. Crude-extract assays confirm glutathione-dependent fosfomycin inactivation under metal-chelating conditions. Distant Gfa homologs also conferred resistance, although activity was restricted to a subset of proteins tested. Together, these findings establish Gfa as a previously unrecognized fosfomycin resistance family and highlight the limitations of homology-based mining for discovering novel resistance determinants and predicting which family members are functional.

## Introduction

Antimicrobial resistance (AMR) represents one of the most pressing threats to global public health, progressively limiting the efficacy of standard antibacterial therapies^1^. The development of novel antibiotic classes has not kept pace with the rapid emergence and dissemination of multidrug-resistant pathogens, leading to a renewed interest in the optimized use of older antibiotics, either alone or in combination with other agents or adjuvants^2^. Among these, colistin and fosfomycin have re-emerged as critical options against multidrug-resistant Gram-negative bacteria^3^. However, the clinical use of colistin has been increasingly restricted due to its well-documented nephrotoxicity and neurotoxicity, which significantly limit dosing and treatment duration; as well as to the global dissemination of plasmid-mediated *mcr* genes that compromise its efficacy^4–6^.

Fosfomycin is a broad-spectrum antibiotic that inhibits MurA (UDP-*N*-acetylglucosamine enolpyruvyl transferase), thereby blocking the first committed step of peptidoglycan biosynthesis^7^. Since its introduction more than five decades ago, fosfomycin has demonstrated sustained clinical efficacy, particularly in uncomplicated urinary tract infections, but also as an intravenous agent and in combination regimens for severe infections caused by multidrug-resistant Enterobacterales and *Pseudomonas aeruginosa*^8–11^. Its favourable pharmacokinetic properties, including high urinary concentrations and the possibility of single-dose oral administration, have contributed to its widespread use as a first-line therapy in several countries and may reduce selection of persistent subpopulations^12,13^. Despite these advantages, the clinical positioning of fosfomycin has recently become more uncertain. Updated susceptibility testing guidelines and breakpoint revisions have highlighted limitations in the predictive value of *in vitro* fosfomycin susceptibility testing, particularly for intravenous use, and have drawn attention to the complexity of interpreting resistance phenotypes^14^. Fosfomycin resistance can arise from chromosomal mutations that reduce antibiotic uptake, particularly those affecting the GlpT and UhpT transport systems or the UhpA regulator^15^. Resistance can also be mediated by fosfomycin-inactivating enzymes, including FosA, FosB and FosX, which use glutathione, bacillithiol and water, respectively, to open the epoxide ring of fosfomycin^16–18^. Interpretation of fosfomycin resistance phenotypes is further complicated by the frequent detection of chromosomal fosfomycin-modifying enzymes in clinically relevant Gram-negative species^19^. Chromosomal *fosA* homologs are widespread among Enterobacteriaceae and *Pseudomonas* spp., and additional fosfomycin-modifying enzymes have recently been identified in other Gram-negative pathogens, including *Morganella* spp. and *Aeromonas* spp.^20,21^. Importantly, the presence of these determinants does not necessarily predict a resistant phenotype, as chromosomal genes may remain weakly expressed or phenotypically silent^22^. Conversely, acquired *fos* genes occur in plasmids, transposons, and integrons^9,23^, where expression may be enhanced by mobile-element-associated promoters^22,23^. Thus, the resistance phenotype associated with a *fos* determinant depends not only on the encoded protein but also on its genetic context and expression level.

Integrons represent highly efficient gene capture and expression platforms that have played a major role in bacterial adaptation to antibiotic pressure and in the dissemination of resistance genes^24,25^. By recruiting cassettes of diverse and often unknown functions, integrons constitute reservoirs of latent adaptive traits that may become clinically relevant under appropriate selective pressures. A substantial proportion of integron cassettes encode proteins of unknown or poorly characterized function, reflecting both their evolutionary diversity and the limited predictive power of sequence-based annotation alone. Resistance genes with no similarity to known determinants may therefore be missed by sequence-based surveillance. In this study, we functionally screened 129 gene cassettes of unknown function (*gcus*) derived from class 1 mobile integrons from a previously established library^26^ -in which bacteriophage defense systems were also identified- to identify novel antimicrobial resistance genes. One gene cassette, *gcu173*, conferred high-level fosfomycin resistance despite lacking detectable homology to any known *fos* gene family. We subsequently undertook a comprehensive genetic, functional, and structural characterization of this determinant, herein referred to as Gfa173. Our results identify Gfa173 as a novel fosfomycin resistance mechanism that differs from canonical Fos enzymes both in its cysteine-rich, metal-associated architecture and in its lack of susceptibility to phosphonoformate inhibition^27,28^. This work expands the known biochemical landscape of fosfomycin resistance and demonstrates that functional screening remains essential to uncover resistance determinants that escape homology-based prediction, with direct implications for surveillance and clinical interpretation.

## Materials and Methods

### Strain collection and growth conditions

A list of bacterial strains used in this study is provided in Table S1. *Escherichia coli* DH5α, MG1655 and BL21 DE3 were used as the cloning, antimicrobial susceptibility testing and protein production hosts, respectively, while *Pseudomonas aeruginosa* PAO1 and *Klebsiella pneumoniae* ATCC 23357 were used as recipients for antimicrobial susceptibility testing only. Strains were grown at 37 °C in Luria Bertani (LB) medium supplemented, when appropriate, with zeocin (50 µg/ml), ampicillin (100 µg/ml), or tetracycline (10 µg/ml) for plasmid maintenance. Liquid cultures were incubated for 16 h in an Infors Multitron shaker at 200 rpm (Infors HT, Switzerland).

### Molecular biology techniques

Gene cassettes were cloned using plasmids specifically designed for functional screening and protein expression. All *gcu* sequences were cloned into the pMBA vector using Gibson assembly as described previously^26,29^. The pMBA vector contains a synthetic region mimicking a class 1 integron platform, including an *attI* site and a disrupted *intI1* gene to prevent undesired recombination events^30^. A *gfp* reporter cassette is located downstream of the *attI* site and acts as a second gene cassette upon *gcu* insertion.

The *gcu173* sequence and selected *gfa*-like homologs were synthesized by Integrated DNA Technologies (IDT, Newark, USA) using eBlocks® gene synthesis and assembled directly into the target plasmid. PCR amplifications were performed using Phusion Green High-Fidelity DNA Polymerase Master Mix (Thermo Fisher Scientific) according to the manufacturer’s instructions. Plasmid constructs were verified by Sanger sequencing performed by Macrogen (Spain) or by long-read plasmid sequencing carried out by Plasmidsaurus (Spain). Site-directed mutagenesis was performed using Gibson assembly with specifically designed primers to introduce targeted amino-acid substitutions (Table S2).

For protein expression, *gfa* and *fos* genes were cloned into the pMAL expression vector under the control of an IPTG-inducible promoter and expressed as His-tagged fusion proteins. For experiments conducted in *P. aeruginosa*, a pMBA-derived plasmid (pBT) carrying a pBBR1 replication origin and a tetracycline resistance marker was used to ensure plasmid replication and maintenance in this host.

### Antimicrobial susceptibility testing

Screening for antimicrobial resistance was performed using Kirby-Bauer disk diffusion method on Mueller-Hinton agar (Oxoid) using antibiotic impregnated disks (Biorad, Spain). The list of antibiotics used is detailed in Table S3.

Fosfomycin disodium salt (TCI, Cymit, Spain) MIC was determined by agar dilution as recommended by the EUCAST guidelines^31^. Glucose-6-phosphate (G6P) (Roche, Germany) was added at a concentration of 25 μg/ml to Mueller-Hinton agar plate with fosfomycin disodium salt concentration ranging from 0.25 to 2,048 μg/ml. Phosphonoformate (PPF) (Sigma Aldrich, Germany) was added at a concentration of 3 mM to assess its inhibitory effect on *fos* or *gfa* genes as described previously^18^. Formaldehyde susceptibility was also determined by agar dilution using Mueller–Hinton agar supplemented with two-fold serial concentrations of formaldehyde. Plates were prepared immediately before inoculation to minimize formaldehyde loss by evaporation^32^.

### *In silico* analysis

We generated the list of *gcu* as described previously using the INTEGRALL database coupled with IntegronFinder^26,33,34^. For *gcu173*, which conferred a fosfomycin resistance phenotype, protein structure prediction was performed using the AlphaFold2 pipeline^35^, followed by Foldseek^36^ to identify potential structural homologs. In parallel, the Iterative Threading ASSEmbly Refinement (I-TASSER)^37^ platform was used to infer putative ligand-binding sites and functional annotations, as AlphaFold2 predictions are limited to apo protein structures and do not provide ligand-binding inference. To investigate the distribution and genetic context of Gfa-like proteins, BLASTp and BLASTn searches were performed against the NCBI non-redundant protein and nucleotide databases to identify homologous sequences and their corresponding genomic environments. The genetic contexts of Gfa homologs were manually inspected to determine their chromosomal or mobile localization. Multiple sequence alignments of Gfa and Gfa-like proteins were generated in Geneious (version 2026.0.2) using the MAFFT algorithm, and phylogenetic analyses were performed based on amino-acid alignments using the neighbour-joining method with default parameters.

### Molecular docking analyses

Molecular docking analyses were performed to investigate the interaction of Gfa with fosfomycin, PPF, and glutathione (GSH). The AlphaFold2-predicted Gfa structure was used as the receptor and prepared using AutoDockTools (version 1.5.7)^38^, including the addition of polar hydrogens and assignment of Gasteiger charges. Ligand structures for fosfomycin, PPF, and GSH were retrieved from public databases or manually built and prepared using Open Babel (version 3.1.1) for geometry optimization and conversion to the appropriate file formats. Docking simulations were carried out using AutoDock Vina (version 1.1.2), with a search grid encompassing the predicted active-site cavity identified from structural analyses. Docking poses were ranked according to Vina scoring functions. Resulting complexes were visualized and analyzed using UCSF ChimeraX (version 1.10.1) to assess ligand orientation, predicted interactions, and consistency with conserved residues and putative metal-binding sites.

### Protein expression, crude extract preparation, and enzymatic assays

Genes encoding Gfa or FosG fosfomycin resistance enzymes were cloned into the pMAL expression vector and transformed into *Escherichia coli* BL21(DE3). Overnight cultures were diluted 1:100 into fresh Luria–Bertani (LB) medium supplemented with the appropriate antibiotic and grown at 37 °C with shaking until mid-exponential phase (OD_600_ ≈ 0.5–0.6). Protein expression was induced by addition of isopropyl β-D-1-thiogalactopyranoside (IPTG) (ThermoFisher, Spain) to a final concentration of 0.5 mM, followed by incubation for 4 h at 37 °C with agitation.

Cells were harvested by centrifugation (5,000 × g, 10 min, 4 °C) and resuspended in lysis buffer consisting of 50 mM Tris-HCl (pH 7.5), 300 mM NaCl, supplemented with phenylmethylsulfonyl fluoride (PMSF) added immediately prior to lysis as a protease inhibitor. Cell disruption was performed by sonication on ice using a probe sonicator with multiple short pulses (typically 10 s ON / 30 s OFF cycles) at 30–40% amplitude to prevent overheating. The total sonication time was adjusted according to pellet size and volume.

Following lysis, cellular debris was removed by centrifugation (4,000 × g, 30 min, 4 °C). The clarified supernatant was subsequently passed through a 0.22 µm filter to obtain sterile crude extracts. Crude extracts were aliquoted and stored at −80 °C until further use. Protein concentration was estimated by absorbance at 280 nm or by Bradford assay when required.

Functional inactivation assays were performed using a disk-based method adapted from the modified carbapenem inactivation method (mCIM)^39^. Briefly, fosfomycin disks (50 μg, Oxoid™, Spain) were incubated for 4 h at 37 °C with crude extracts diluted either in Tris buffer (5 mM) supplemented with GSH (10 μM), NaCl (34 mM) and KCl (22 mM) at pH 7.4 or Mueller–Hinton (MH) broth in the presence of GSH (10 μM), ZnCl_2_ (10 μM), MnCl_2_ (1 mM), PPF (3 mM) or EDTA (10 mM) alone or in combination. After incubation, disks were placed onto MH agar plates inoculated with a fosfomycin-susceptible *E. coli* indicator strain. Inhibition zones were measured after overnight incubation and compared with appropriate negative controls (buffer alone or crude extract from empty-vector strains). These assays were used to qualitatively assess fosfomycin inactivation by Gfa or Fos-containing extracts under different biochemical conditions.

### *attC* recombination experiments

*attC* recombination assays were performed as previously described^40^, with minor adaptations. Briefly, the 3′ region of the *gcu173* cassette, encompassing its *attC* site, was cloned into the mobilizable plasmid pSW23T —in both orientations to compare top and bottom strand recombinations— carrying a chloramphenicol resistance marker. The resulting construct was introduced into *E. coli* β2163 (*dapA, pir*^+^), which requires diaminopimelic acid (DAP) supplementation for growth.

*E. coli* β2163 harbouring the pSW23T-*gcu173* construct was used as the donor strain in conjugation experiments with *E. coli* DH5α as the recipient strain. The recipient carried plasmid p3938 (Carb^R^) encoding an L-arabinose–inducible *intI1* gene and plasmid p929 (Kan^R^) containing a single attI1 site. Conjugation experiments were performed on LB agar plates supplemented with DAP and L-arabinose (0.2%) and incubated overnight at 37 °C.

Following cell recovery, ten-fold serial dilutions were plated onto LB agar supplemented with carbenicillin (100 µg/ml) and glucose (1%) to select for the total recipient population, or with carbenicillin, glucose, and chloramphenicol (25 µg/ml) to selectively recover recombination events resulting from *attC* × *attI* integration.

### Fitness cost experiments

To assess the relative fitness cost associated with *gcu173* expression and to compare it with other fosfomycin resistance determinants, in vitro competition assays were performed following a previously described protocol with minor modifications^41^.

For initial benchmarking, *gcu173* and selected *fos* determinants were cloned into the pMBA expression vector with the *gfp* gene carrying two stop codon mutations at the beginning of its sequence (pMBA-*Δgfp*) and competed against an *E. coli* MG1655 strain carrying the pMBA-GFP-WT plasmid as a reference control.

To enable direct comparison among Gfa, Gfa-like homologs and FosG under identical conditions, *gfa173* and selected *gfa*-like genes were cloned into the modified pMBA-*Δgfp* plasmid. These GFP-negative strains were competed against a common *E. coli* MG1655 reference strain carrying a functional *gfp* gene integrated chromosomally. This design allows comparative assessment of relative fitness costs across Gfa-like proteins using a single standardized competitor.

Six independent biological replicates of each construct were grown overnight at 37 °C in 96-well plates containing LB broth supplemented with zeocin (50 µg/ml). Overnight cultures were mixed at a 1:1 volumetric ratio between GFP-negative (pMBA*-Δgfp*) strains and the GFP-positive reference strain. Mixed cultures were diluted 1:20 into 96-well plates containing 0.9% NaCl and subsequently diluted 1:20 into two additional plates, one containing 0.9% NaCl and the other containing LB broth supplemented with zeocin. The first plate was immediately used to determine the relative proportions of GFP-positive and GFP-negative cells at day 1 using a CytoFLEX S flow cytometer (Beckman Coulter, USA). The second plate was incubated overnight at 37 °C. On the following day, cultures were diluted 1:400 in 0.9% NaCl and analysed again by flow cytometry to determine strain proportions at day 2. Relative fitness cost was calculated as the change in GFP fluorescence ratios between day 1 and day 2.

### Statistical analysis of fitness effects

Fitness effects were statistically assessed using a linear regression model, following the approach described by Hipólito et al.^42^. Relative fitness was used as the response variable and the resistance gene as a categorical explanatory variable. Gene-associated fitness effects were classified as costly, neutral, or beneficial according to whether the estimated effect was significantly below, not significantly different from, or significantly above the neutral fitness reference, respectively. Statistical significance was defined as *p* < 0.05. Six independent biological replicates were included for each construct.

## RESULTS

### Gcu173 is a novel Gfa-like fosfomycin resistance determinant already disseminated in pathogens

To identify novel antimicrobial resistance determinants encoded by integron gene cassettes, we functionally screened a collection of 129 gene cassettes cloned into the pMBA vector and assessed their impact on antibiotic susceptibility using disk diffusion assays. Among these, one cassette, *gcu173*, reproducibly conferred resistance to fosfomycin when expressed in *E. coli* DH5α. The *gcu173* gene is 435 nucleotides in length and encodes a 144–amino-acid protein. Sequence analysis revealed that *gcu173* shared no detectable homology with any previously described *fos* gene family. Structural homology searches, however, identified similarity to a gene annotated as *gfa* (<u>g</u>lutathione-dependent formaldehyde-<u>a</u>ctivating gene), and we therefore adopted the designation Gfa for this determinant while conserving the original number of the *gcu*. Hence we will refer to *gfa173* and Gfa173 for this specific gene and encoded protein, respectively.

Database searches subsequently identified several close Gfa homologs (>90% amino-acid identity) in diverse bacterial species, including clinically relevant taxa such as *Pseudomonas putida* and *Providencia rettgeri*. In *P. putida* and a Pseudomonadaceae bacterium, these homologs were embedded within canonical class 1 integron structures, suggesting recent mobilization (Figure 1A). In contrast, the *gfa* homolog identified in *Providencia rettgeri* was associated with a chromosomal integron detected by IntegronFinder and carrying a different integrase class. This integrase shared only 42% and 54.2% amino-acid identity with IntI1 and IntI2, respectively, indicating that *gfa173*-like can also be found in sedentary chromosomal integrons (Figure 1A). Given their high sequence conservation and conserved genetic organization, these close homologs likely represent functionally equivalent variants of Gfa. Their presence within mobile gene capture platforms in opportunistic pathogens indicates that Gfa-like determinants are already present in clinically relevant bacterial populations rather than confined to environmental reservoirs.

**Figure 1.**
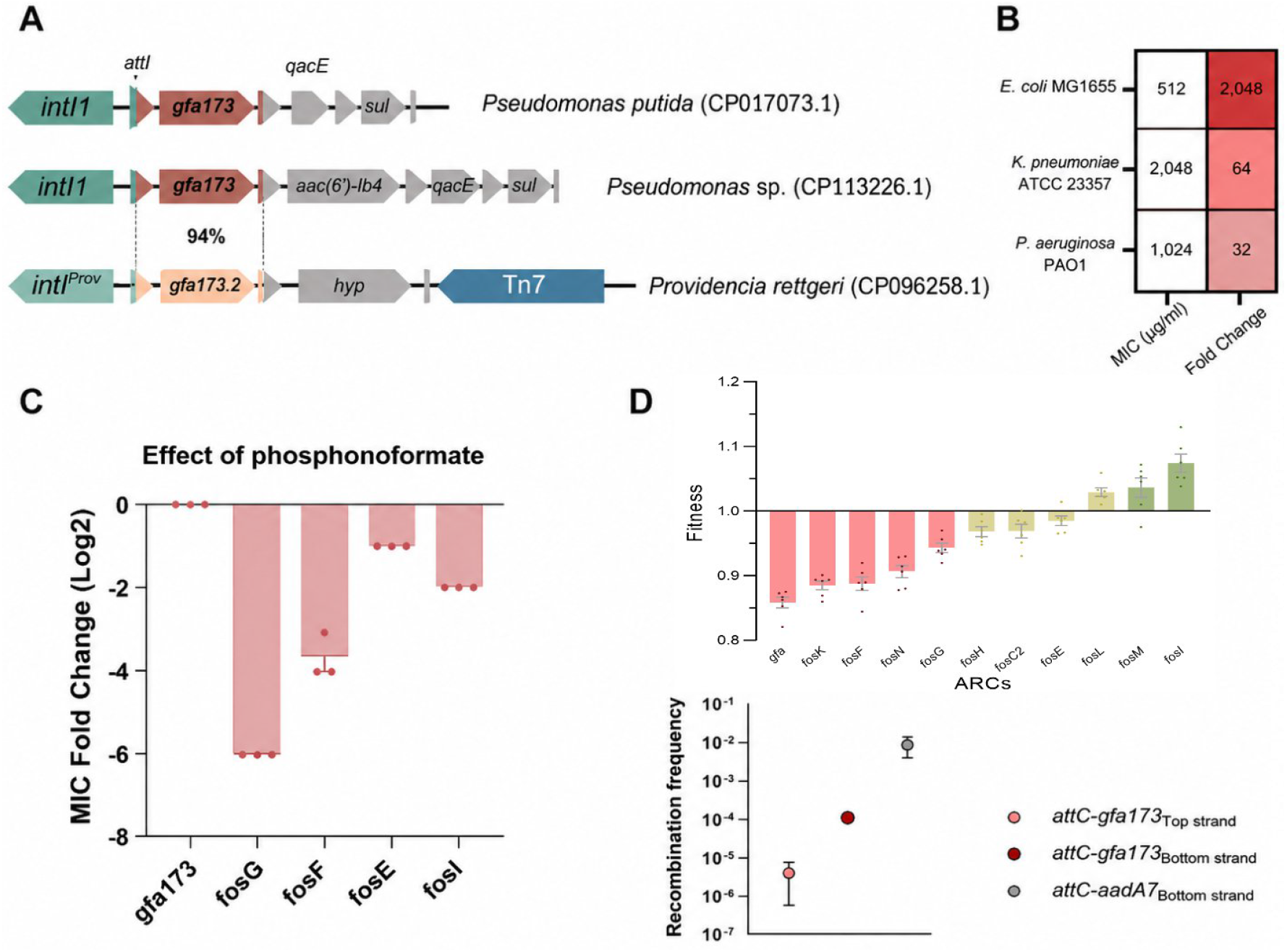
**(A)** Genetic contexts of *gfa173* and closely related *gfa173* homologs identified in bacterial genomes. In *Pseudomonas putida* and a Pseudomonadaceae bacterium, *gfa173* is embedded within canonical class 1 integron platforms downstream of *intI1*, together with other resistance gene cassettes. In *Providencia rettgeri*, the *gfa173* homolog (*gfa173*.*2*) is associated with a sedentary chromosomal integron carrying a divergent integrase (*intI*^*Prov*^) and is linked to the Tn*7* transposon. Percent amino-acid identity between homologs is indicated where applicable. Accession numbers are shown in parentheses. **(B)** Fold change in fosfomycin minimum inhibitory concentration conferred by *gfa173* expression in different bacterial hosts. Values represent the increase in MIC relative to the corresponding control strain for *E. coli* MG1655, *K. pneumoniae* ATCC 23357, and *P. aeruginosa* PAO1. Expression of *gfa173* resulted in 2,048-fold, 64-fold, and 32-fold increases in fosfomycin MIC, respectively. **(C)** Effect of phosphonoformate (PPF) on fosfomycin resistance mediated by integron-borne *fos* genes or by *gfa173*. Fold changes in fosfomycin MICs (log_2_ scale) were determined in the presence of 3 mM of PPF. **(D)** Fitness cost and recombination capacity of integron-associated fosfomycin resistance determinants. Top: relative fitness values measured by pairwise competition assays in *E. coli* MG1655 colours indicate whether the gene is significantly costly (red), neutral (yellow) or confers a fitness gain (green) using the linear regression approach described by Hipólito et al^42^. Each dot represents one biological replicate; bars indicate mean values. Bottom: recombination frequencies of *attC* sites associated with *gfa173* and *aadA7* control cassettes, measured for top and bottom strands in an IntI1-dependent recombination assay.

Agar dilution assays revealed that expression of *gfa173* conferred a fosfomycin MIC of 512 µg/mL, corresponding to a >2,000-fold increase compared with the parental *E. coli* MG1655 strain (MIC = 0.25 µg/mL). To evaluate whether Gfa-mediated resistance was conserved across species, *gcu173* was introduced into *K. pneumoniae* ATCC 23357 using pMBA and into *P. aeruginosa* PAO1 using pBT, a pMBA-derived plasmid carrying the pBBR1 replicon compatible with *Pseudomonas* spp. In both hosts, *gfa173* expression resulted in marked increases in fosfomycin MICs (2,048 µg/mL in *K. pneumoniae* and 1,024 µg/mL in *P. aeruginosa*), representing 64-fold and 32-fold increases relative to the corresponding control strains (Figure 1B). Despite its annotation as a glutathione-dependent formaldehyde-activating enzyme, expression of *gfa173* did not alter formaldehyde susceptibility compared with the empty-vector control (MIC = 19.53 mg/L for both strains) (Table S5).

### Gfa173 is not sensitive to phosphonoformate inhibition

One characteristic shared by most enzymatic fosfomycin resistance determinants is their susceptibility to inhibition by phosphonoformate (PPF, foscarnet®). PPF acts as a transition-state analogue inhibitor and efficiently blocks thiol-dependent fosfomycin resistance enzymes, including the glutathione-S-transferase (FosA) and bacillithiol-S-transferase (FosB) families, by coordinating the divalent metal ion in the catalytic site and preventing fosfomycin modification^27,28^. As a result, PPF has been proposed as a potential adjuvant to restore fosfomycin activity against bacteria expressing FosA- or FosB-mediated resistance. In contrast, members of the FosX family, which are metalloenzymes that hydrolyse fosfomycin rather than conjugating it to a thiol, have been shown to be intrinsically resistant to phosphonoformate inhibition, displaying little or no reduction in fosfomycin MIC in the presence of PPF^18,43^.

To assess the susceptibility of Gfa to phosphonoformate inhibition, fosfomycin MICs were determined by agar dilution on MH agar in the presence or absence of 3 mM PPF (Figure 1C). Four integron-borne *fos* genes were included as controls: *fosG* and *fosF*, encoding glutathione-dependent transferases, and *fosE* and *fosI*, encoding FosX-like enzymes. In the presence of PPF, the MICs of clones carrying *fosG* and *fosF* decreased by 64-fold and 8-fold, respectively, whereas clones carrying *fosE* and *fosI* showed only modest reductions of 2-fold and 4-fold. In contrast, no difference in fosfomycin MIC was observed for the clone carrying *gfa* in the presence or absence of PPF, indicating that Gfa-mediated fosfomycin resistance is not inhibited by PPF (Figure 1C).

### *gfa173* imposes a measurable fitness cost and is mobilized by integrons

To understand how likely is Gfa to subsist in populations in the absence of antibiotics, we determined the fitness effects of Gfa through competition assays and compared it to those of other fosfomycin resistance determinants in integrons. Results revealed substantial heterogeneity in the fitness effects imposed by integron-borne fosfomycin resistance determinants (Figure 1D). While several *fos* genes exhibited neutral or even mildly beneficial effects on host fitness, others imposed a variety of costs in the absence of the antibiotic. Within this spectrum, expression of *gfa173* resulted in the strongest reduction in relative fitness among all determinants tested, placing it at the extreme low end of the fitness distribution. This contrasts with most of other integron-associated *fos* genes, which clustered closer to neutrality and, in some cases, conferred a slight fitness advantage under the same experimental conditions (Figure 1D).

### Gfa173 is a *bona fide* integron cassette with moderate mobility

*gfa173* exhibits all the structural hallmarks of a functional integron cassette. *In silico* analysis of the associated *attC* site revealed the well-defined secondary structure ^44^ with a canonical L box and R box, as well as an extrahelical base, consistent with efficient recognition by integron integrases^45^. The predicted folding of the *attC* site was highly stable (ΔG = −26.37 kcal/mol), supporting its recombination ability (Figure S1). To confirm this, we performed functional recombination assays that confirmed that the *gfa173 attC* site is recombined by IntI1. The bottom strand showed higher recombination frequencies (1.09 × 10^−4^) than the top strand (2.58 × 10^−6^), in line with the known strand bias in integron recombination (Figure 1D)^45^. The frequency of bottom strand recombination is limited compared to the *attC* of *aadA7* (∼10^-2^), used here as a highly recombinogenic control.

Together, these results demonstrate that *gfa173* is a *bona fide* integron gene cassette and suggest that its relatively low recombination frequency and substantial fitness cost may represent two barriers to its dissemination.

### Structural modelling of Gfa173 reveals a conserved cysteine-rich, metal-associated architecture

Structure prediction using the AlphaFold2 confidently suggests that Gfa173 forms a homodimer (ipTM = 0.93; pTM = 0.93); with the ipTM score reflecting confidence in the relative positioning of the two chains and the pTM score reflecting confidence in the overall predicted structure (Figure 2A).

**Figure 2.**
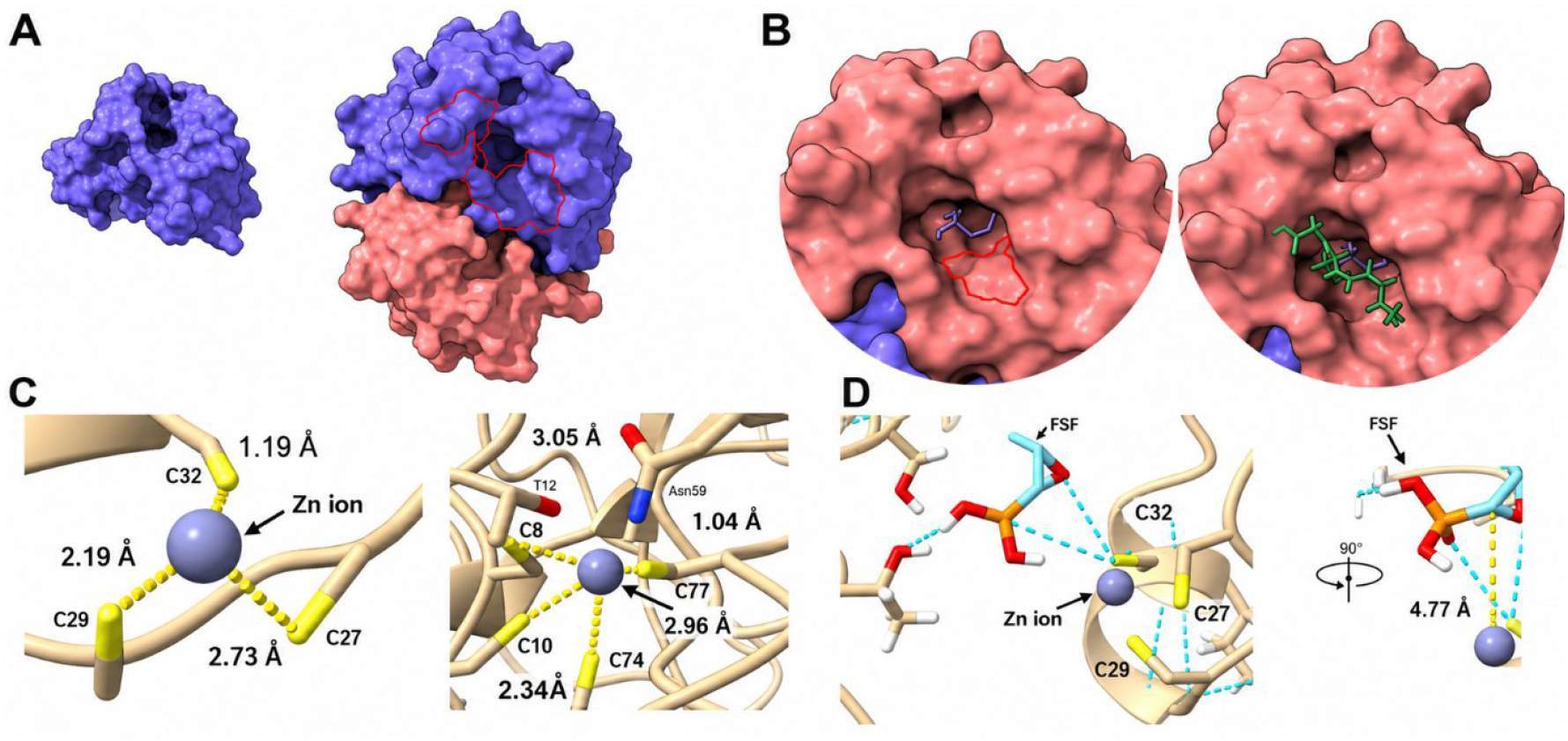
**(A)** Predicted three-dimensional structure of Gfa173. Left, surface representation of the Gfa173 monomer. Right, homodimeric assembly predicted by AlphaFold2, visualized using ChimeraX. The two protomers are shown in blue and salmon. The predicted glutathione-binding region, encompassing the first conserved motif, is highlighted in red on the dimeric structure. **(B)** Molecular docking of fosfomycin and glutathione within the Gfa173 active-site cavity. Left, docking of fosfomycin alone reveals its positioning within a surface-accessible pocket of Gfa173 and proximity to the cysteine-rich region C27-C29-C32 highlighted in red. Right, docking of fosfomycin followed by glutathione shows simultaneous accommodation of both ligands within the cavity, consistent with an ordered or substrate-dependent binding mode. **(C)** Predicted structural organization of the two cysteine-rich zinc-binding sites identified in Gfa173. Close-up views show zinc coordination by conserved cysteine residues, with metal–sulfur distances indicated. The zinc-binding site formed by C27, C29, and C32 is shown on the left, whereas the second zinc-binding site involving C8, C10, C74, and C77 is shown on the right. **(D)** Detailed view of the zinc-binding site overlapping with the ligand-binding cavity. Zinc coordination by C27, C29, and C32 constrains the orientation of the cysteine-rich scaffold. Docking of fosfomycin in the zinc-bound structure reveals a hydrogen bond between fosfomycin and a zinc-coordinated cysteine residue, indicating that zinc indirectly enables antibiotic binding by organizing the local cysteine architecture rather than by direct metal–ligand interaction.

To investigate potential functional features, Gfa173 was further analysed using the I-TASSER platform. This analysis identified two putative zinc-binding sites and one predicted glutathione-binding site. The latter comprises residues C29, S30, K31, E67, N68, V69, K71, F112, and S115. The first predicted zinc-binding site is formed by residues C8, C10, C74, and C77, whereas the second involves residues C27, C29, and C32, revealing a prominent cysteine-rich architecture consistent with metal coordination^46^.

Structural similarity searches performed using Foldseek against the PDB100 database identified PA2722 from *Pseudomonas aeruginosa* (PDB: 8AJQ), a glutathione-dependent formaldehyde-activating enzyme, as the closest structural homolog of Gfa173 when the predicted homodimer was used as query. Despite low sequence identity (∼20%), high structural similarity scores were obtained (qTM ≈ 0.78; tTM ≈ 0.85); where qTM represents query-normalized and tTM target-normalized similarity. Complementary Foldseek searches using the Gfa173 monomer against the AlphaFold database retrieved numerous proteins annotated as Gfa or aldehyde-activating enzymes across diverse Proteobacteria, with higher sequence identity (35–60%), showing the widespread distribution of Gfa-like proteins among different bacterial species.

To explore the binding of Gfa173 to its putative substrates and cofactors, we performed molecular docking analyses. Docking simulations showed that GSH, fosfomycin, and formaldehyde could each be accommodated within the predicted catalytic cavity of Gfa173 with consistent and convergent binding poses. Notably, ligand binding was order-dependent. When glutathione was docked first, subsequent docking of fosfomycin was not possible, suggesting steric or conformational exclusion. In contrast, docking fosfomycin prior to glutathione allowed simultaneous accommodation of both ligands within the active site (Figure 2B). In this configuration, a hydrogen bond was consistently predicted between the phosphonate group of fosfomycin and a cysteine residue located within a compact zinc-coordinated cluster. Visualization of metal–sulfur distances and coordination geometry indicated stable zinc coordination by multiple conserved cysteine residues, consistent with a structural rather than catalytic metal-binding role (Figure 2C). Notably, this spatial overlap is restricted to the zinc-binding site formed by C27, C29, and C32, whereas the second predicted zinc-binding site is located in a distinct region of the protein. To further assess the influence of zinc coordination on ligand positioning, we performed docking analyses using the zinc-bound Gfa173 structure. In this context, zinc coordination constrained the relative orientation of cysteine residues, resulting in a stable configuration in which fosfomycin maintained a hydrogen bond with a zinc-coordinated cysteine residue (Figure 2D). These observations suggest that zinc does not directly interact with fosfomycin but instead organizes a cysteine-rich scaffold that enables antibiotic binding. Together, these results support a model in which Gfa173 relies on a conserved cysteine-and metal-organized structural framework to accommodate fosfomycin and glutathione.

### Conserved cysteines are required for Gfa173 activity

To confirm the contribution of residues located within or close to the predicted ligand-binding region, alanine substitutions were introduced based on the I-TASSER and docking predictions (C29A, S30A, S66A, N68A, T70A, K71A, S115A, K116A, and R140A). With the exception of C29A, all these substitutions resulted in partial but reproducible reductions in fosfomycin MICs, indicating that these residues contribute to, but are not strictly required for, the resistance phenotype (Figure 3A).

**Figure 3.**
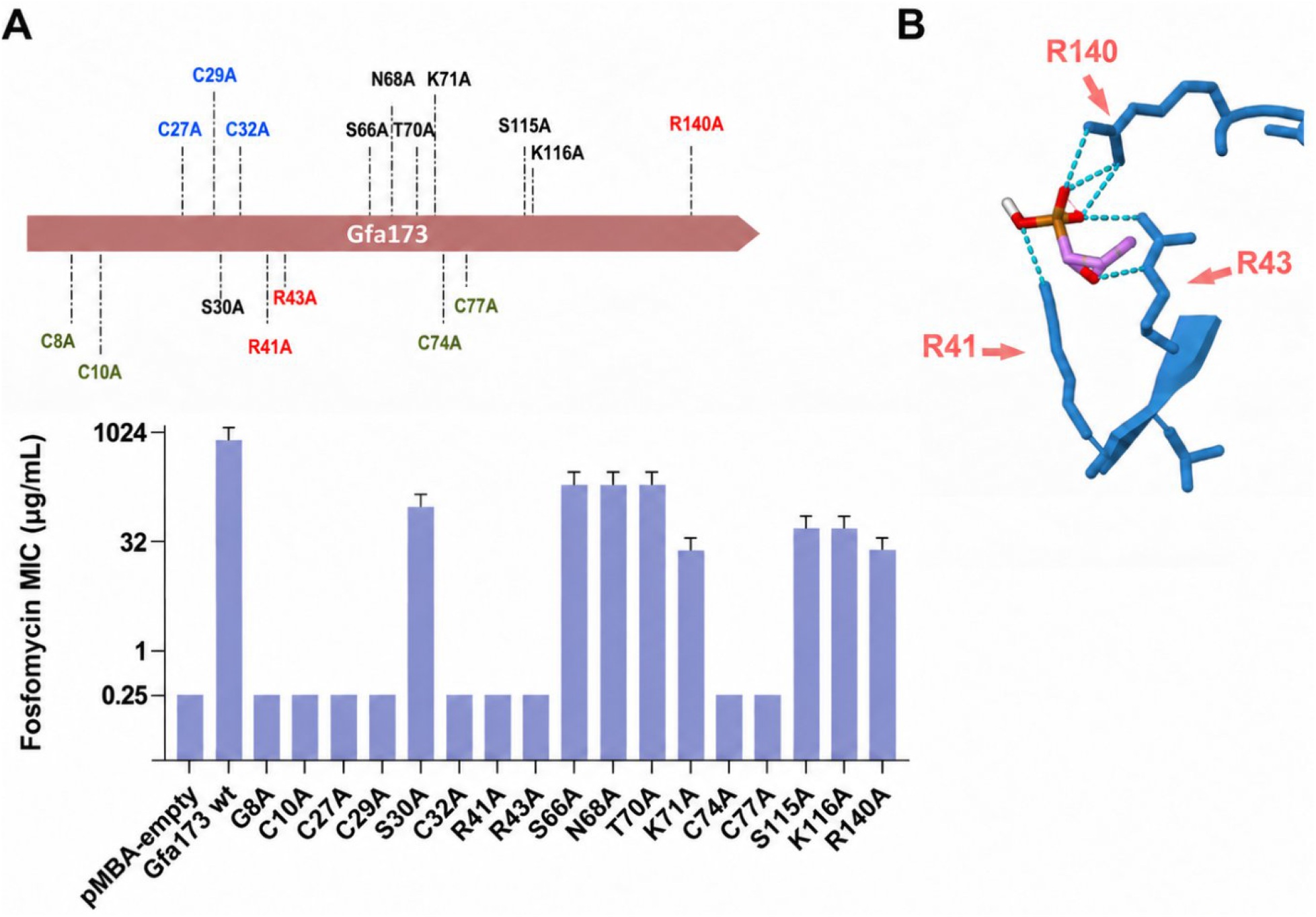
(A) Schematic representation of the Gfa173 protein indicating the positions of residues targeted by alanine substitution (top) and corresponding fosfomycin minimum inhibitory concentrations (MICs) determined for each mutant (bottom) by agar dilution in duplicate. Bars represent mean MIC values and error bars indicate standard deviations. Residues are color-coded according to their predicted structural or functional role. Cysteines belonging to the zinc-binding cluster overlapping with the predicted ligand-binding pocket (C27, C29, C32) are shown in blue. Cysteines forming a second, spatially distinct cysteine-rich cluster (C8, C10, C74, C77) are shown in green. Arginine residues identified in the docking analysis (R41, R43, R140) are highlighted in pink. R41A and R43A abolished resistance, whereas R140A resulted in a partial MIC reduction. Other residues whose substitution resulted in partial MIC reductions (S30, S66, N68, T70, K71, S115, K116) are shown in black. (B) Structural model illustrating the predicted interaction of fosfomycin within the Gfa173 binding pocket, highlighting R41, R43, and R140 as residues predicted to interact with the substrate.

In contrast, substitution of the conserved cysteine residue C29, which is located within the predicted ligand-binding pocket and also participates in one of the predicted zinc-binding motifs, completely abolished fosfomycin resistance, reducing MIC values to those of the empty-vector control. On this basis, we systematically performed alanine substitutions of all cysteine residues predicted to contribute to metal binding or structural integrity. Strikingly, alanine substitution of any cysteine residue resulted in complete loss of fosfomycin resistance, irrespective of its position within the predicted zinc-binding sites or elsewhere in the protein. In addition, R41A and R43A completely abolished the resistance phenotype. Structural modelling predicts that R41, R43, and R140 can interact with fosfomycin within the predicted binding pocket; however, whereas substitution of R41 or R43 abolished resistance, R140A retained partial activity. Altogether, these data suggest that both the cysteine-rich architecture and specific substrate-interacting residues are important for Gfa173-mediated fosfomycin resistance.

### Gfa173 homologs are widespread and partially functional

To assess whether fosfomycin resistance is conserved across the Gfa family, we selected a representative set of homologs spanning different genetic contexts and levels of sequence identity to Gfa173 (Figure 4A-B). Two Gfa-like proteins, Gfa^CellV1^ and Gfa^CellV2^, were identified adjacent within the same *Cellvibrio* sp. genome and showed 86.1% and 31.9% amino-acid identity with Gfa173, respectively. A third *Cellvibrio*-associated homolog, Gfa^KYGH^, originated from a distinct genomic background and displayed 75.2% identity to Gfa173. *Marinobacter fuscus* genome contained Gfa^fuscus^ sharing 69.4% identity and was located within a transposon-associated region. Finally, we found Gfa^PAO1^ from *P. aeruginosa* PAO1, representing the most divergent homolog examined (20.8% identity). Together, this panel encompassed closely related and divergent Gfa-like proteins from distinct chromosomal and mobile genetic contexts. All *gfa*-like genes were successfully cloned under the weak constitutive PcW promoter. However, repeated attempts to generate PcS-driven constructs for several homologs were unsuccessful, suggesting that stronger expression may impose a substantial burden on the host. Fosfomycin susceptibility testing revealed pronounced functional heterogeneity among Gfa-like proteins (Figure 4D). Under standardized PcW-driven expression in *E. coli, gfa173* conferred a fosfomycin MIC of 32 µg/mL (versus 512 µg/mL with the PcS) . Two additional homologs displayed partial activity, with *gfa*^CellV1^ and *gfa*^KYGH^ conferring MICs of 8/16 µg/mL and 4/8 µg/mL, respectively. In contrast, *gfa*^CellV2^, *gfa*^fuscus^, and *gfa*^PAO1^ failed to increase fosfomycin MIC above baseline levels (<0.5 µg/mL), despite identical expression conditions (Figure 4D). To assess potential host-specific effects, selected homologs were additionally expressed in *K. pneumoniae* and *P. aeruginosa* PAO1. In both alternative genetic backgrounds, fosfomycin susceptibility profiles were fully consistent with those obtained in *E. coli*, confirming that resistance activity is independent of host context and instead reflects intrinsic functional properties of the proteins. Notably, the endogenous Gfa^PAO1^ homolog remained inactive in both native and heterologous hosts, indicating that it is not a fosfomycin resistance determinant even in its original host.

**Figure 4.**
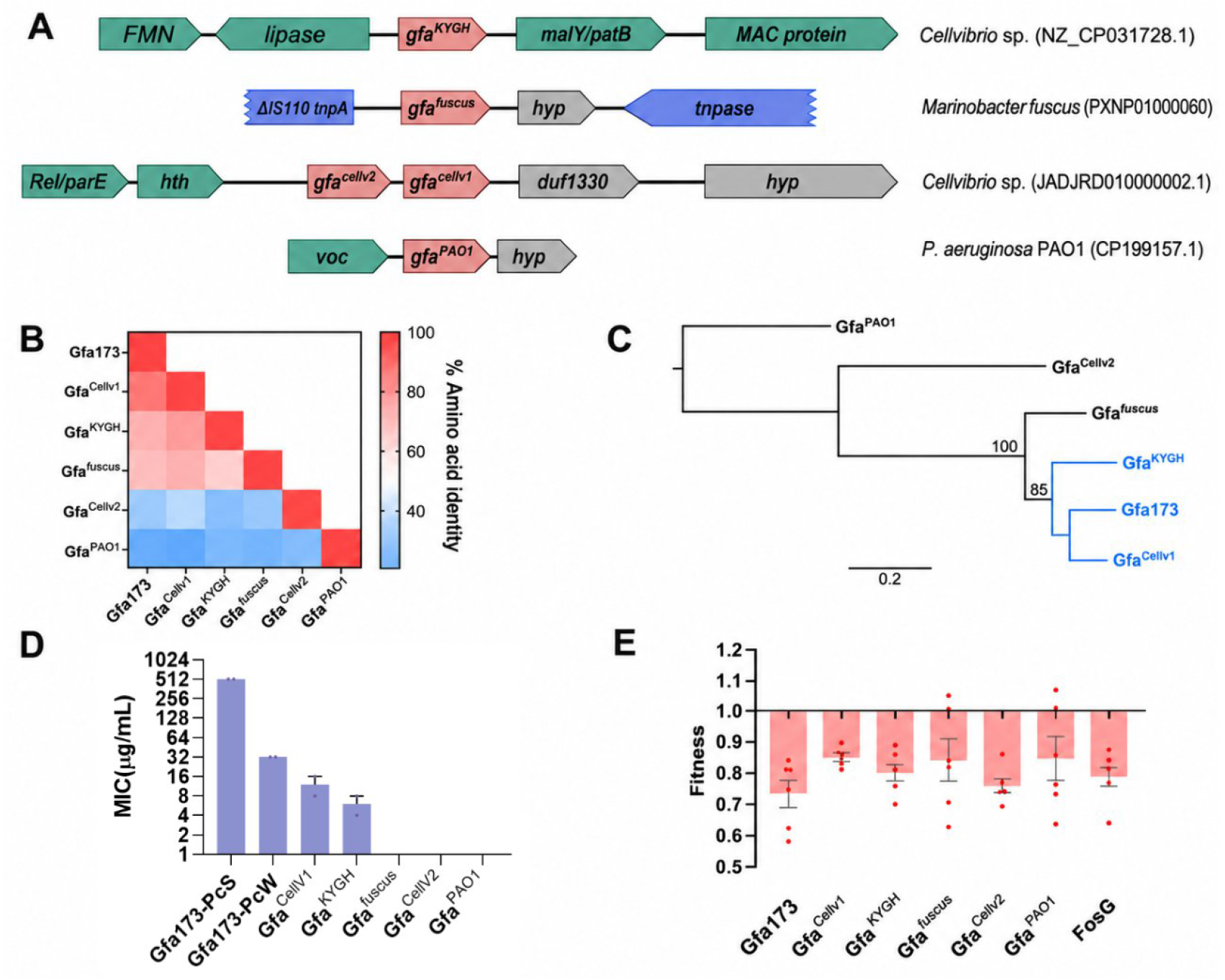
(A) Genetic contexts of representative *gfa*-like distant homologs. Schematic representation of the genomic environments surrounding selected *gfa*-like genes from diverse bacterial species. Functional homologs (*gfa173, gfa*^CellV1^, *gfa*^KYGH^) and non-functional homologs (*gfa*^fuscus^, *gfa*^PAO1^, *gfa*^CellV2^) are shown in their native chromosomal contexts. Genes are colored by functional category, with *gfa*-like genes highlighted in red, mobile elements in blue, genes involved in metabolism in green, and genes encoding hypothetical proteins in grey. (B) Pairwise amino acid sequence identity among Gfa-like homologs. Heatmap showing percentage amino-acid identity between Gfa173 and the tested homologs. Functional proteins exhibit higher sequence identity to Gfa173, whereas non-functional homologs display lower identity values. (C) Phylogenetic reconstruction of Gfa-like proteins. Neighbour-joining phylogenetic tree based on protein sequence alignment of the analyzed Gfa-like homologs. Functional homologs are highlighted in blue. (D) Fosfomycin resistance conferred by Gfa-like homologs. Minimum inhibitory concentrations (MICs) of fosfomycin measured in *Escherichia coli* expressing individual *gfa*-like genes. (E) Fitness cost associated with expression of Gfa-like proteins. Relative fitness measured by pairwise competition assays for strains expressing functional and non-functional Gfa-like homologs, compared with a control strain and FosG.

Phylogenetic reconstruction based on protein sequence alignment revealed a clear subdivision within the Gfa-like family (Figure 4C). The three functional homologs—Gfa173, Gfa^CellV1^, and Gfa^KYGH^—clustered within the same subclade, whereas non-functional homologs formed more divergent branches. Pairwise sequence identity relative to Gfa173 ranged from approximately 20% to 86%, with proteins active on fosfomycin consistently exhibiting higher identity values (≥75%) (Figure 4B). Despite this segregation, multiple sequence alignment demonstrated strict conservation of several cysteine residues across all homologs, including those lacking a resistance phenotype (Figure S2). Consistent with this observation, structural superposition of AlphaFold-predicted models showed that all homologs adopt highly similar three-dimensional folds, with low RMSD values (<1.1 Å) upon alignment to Gfa173, irrespective of phenotype (Figure S2). Thus, neither global sequence similarity nor overall protein architecture discriminates resistance-conferring from non-resistance-conferring Gfa-like proteins, suggesting that fosfomycin resistance depends on a combination of sequence features that cannot be attributed to a single conserved residue or structural motif. To further test the basis of functional divergence within the Gfa family, *gfa*^fuscus^ was selected for targeted mutagenesis, as it occupies an intermediate position between functional and non-functional homologs in terms of sequence identity and structural similarity. The expression of *gfa*^fuscus^ did not result in increased fosfomycin MIC, despite sharing the conserved cysteine-rich, metal-associated architecture characteristic of Gfa173 and having amino acid identity levels comparable to others that do confer resistance. A stepwise mutational strategy was therefore implemented to test if resistance could arise through limited residue replacement. Residues consistently differing between functional (Gfa173, Gfa^CellV1^, Gfa^KYGH^) and non-functional homologs were prioritized, and the substitutions V4M, F25Y, S48K, D49N, Y72F, TR61/62SS, G64K, DR87/88NK, T113V, and SR137/138EA were introduced individually and in combination (Figure S3). None of the resulting variants conferred measurable fosfomycin resistance, with MIC values remaining at baseline levels (0.25 µg/mL). These results indicate that restoration of resistance cannot be achieved through limited residue replacement and that functional divergence of *gfa*^fuscus^ is not governed by a small number of catalytic or binding-site substitutions.

Finally, fitness effects associated with representative functional and non-functional Gfa-like homologs were assessed using pairwise competition assays under standardized PcW-driven expression conditions (Figure 4E). All tested Gfa-like proteins showed relative fitness values below 1, ranging from approximately 0.73 to 0.85. Among the homologs tested, Gfa173 displayed the lowest relative fitness. Importantly, fitness cost did not correlate with fosfomycin resistance phenotype, as several non-functional homologs—including *gfa*^fuscus^ and *gfa*^*cellV2*^ — also displayed a significant cost. Together, these results demonstrate that fosfomycin resistance is not a universal property of Gfa-like proteins. Although the overall structure, cysteine-rich architecture, and predicted metal- and glutathione-binding features are broadly conserved across Gfa-like proteins, fosfomycin resistance is restricted to a subset of the homologs tested.

### Crude extract assays support enzymatic inactivation and define ion dependence

To obtain functional evidence of fosfomycin inactivation by the Gfa protein, we implemented a modified mCIM-like disk inactivation assay using crude cell extracts. In this assay, fosfomycin disks are pre-incubated for 4 h with crude extracts of a strain expressing *gfa173* or *fosG* (as a control) in small volumes of defined incubation mixtures (Mueller–Hinton (MH) broth or Tris buffer) supplemented with selected additives like GSH, divalent ions, PPF, or EDTA. We subsequently used this disk to perform diffusion antibiograms, placing them onto a lawn of a fosfomycin-susceptible strain to quantify residual activity. Under these conditions, Gfa173-mediated fosfomycin inactivation was consistently observed when glutathione was included and disk incubation was performed in MH broth. Replacement of MH by a Tris-based buffer supplemented with glutathione resulted in a weaker and less reproducible inactivation. Complete inactivation under Tris conditions was observed only when EDTA was additionally included, whereas supplementation with Zn^2+^ did not restore MH-like activity. These findings establish that glutathione is required for Gfa173-mediated fosfomycin inactivation and demonstrate that EDTA restores full activity under defined buffer conditions, while exogenous Zn^2+^ does not (Figure 5).

**Figure 5.**
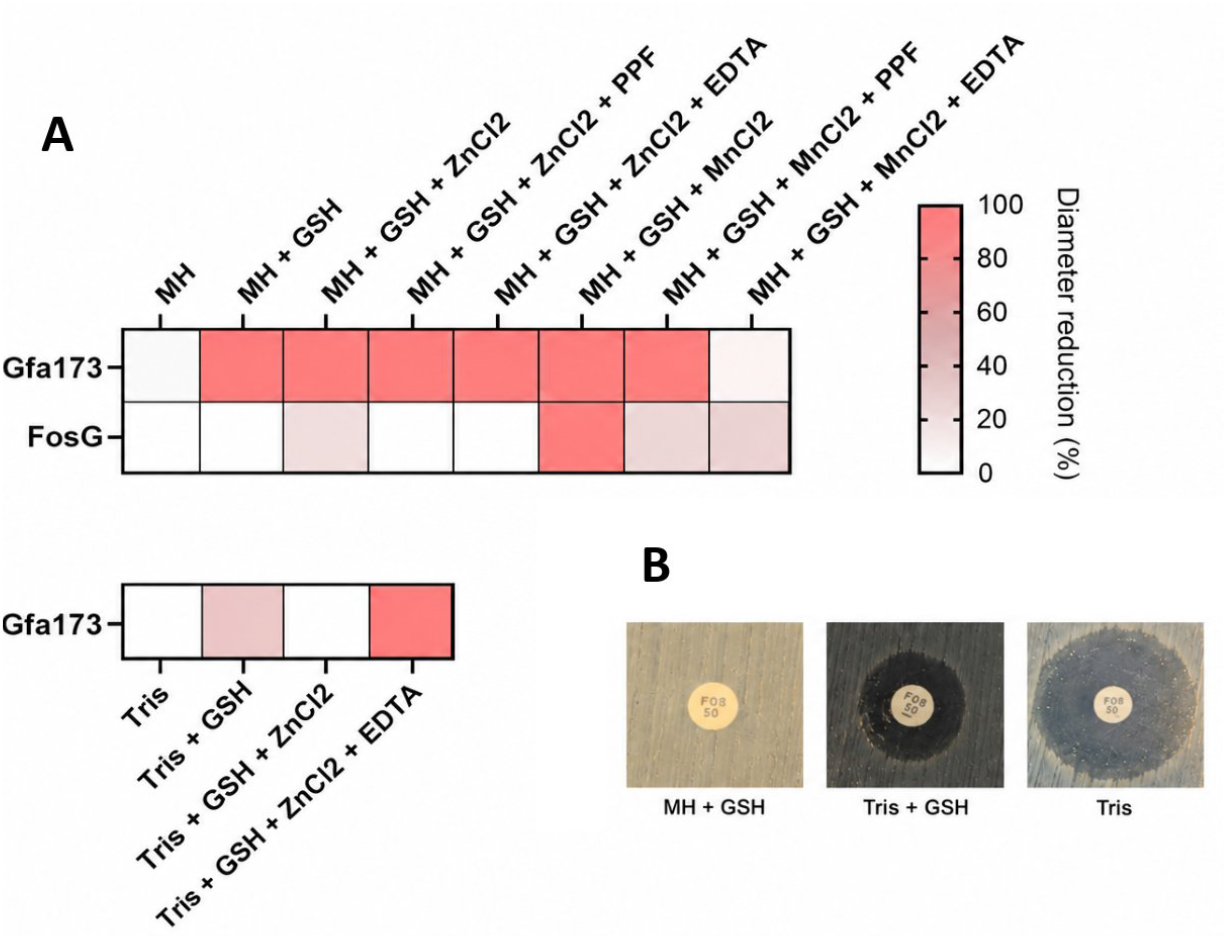
Gfa173-mediated fosfomycin inactivation under different biochemical conditions. **(A)** Fosfomycin disks (50 µg) were pre-incubated for 4 h at 37 °C with crude cell extracts producing Gfa173 or FosG under the indicated conditions and subsequently placed on Mueller–Hinton (MH) agar seeded with a fosfomycin-susceptible *E. coli* indicator strain. Top, assays performed in MH broth supplemented, as indicated, with glutathione (GSH, 10 µM), ZnCl_2_ (10 µM), MnCl_2_ (1 mM), phosphonoformate (PPF, 3 mM), and/or EDTA (10 mM). Bottom, Gfa173 activity assessed in Tris-based buffer under the indicated conditions. Heatmaps represent the percentage reduction in inhibition-zone diameter relative to the corresponding negative control. **(B)** Representative inhibition zones obtained after incubation of fosfomycin disks with Gfa173 crude extracts under MH + GSH, Tris + GSH, and Tris conditions.

Crude extracts expressing FosG displayed a distinct cofactor profile, dependent on both glutathione and Mn^2+^, consistent with a metal-dependent GSH transferase-like activity. FosG activity was strongly reduced by PPF and EDTA, while Gfa remained active in the presence of both compounds, consistent with our previous observations, and indicating that chelation does not suppress Gfa activity under crude-extract assay conditions.

Together, these findings demonstrate that Gfa173 is mechanistically distinct from canonical Fos enzymes. Unlike FosA-like proteins, Gfa173 does not appear to require a freely exchangeable divalent metal cofactor for fosfomycin inactivation and remains active under metal-chelating conditions. Furthermore, Gfa173 retains activity in the presence of PPF, confirming its resistance to a Fos inhibitor and reinforcing its mechanistic divergence. Interestingly, the increased activity observed in the presence of EDTA under Tris-buffered conditions suggests that freely available metal ions may interfere with Gfa173 activity rather than act as obligatory catalytic cofactors; possible explanations for this effect are discussed below.

## DISCUSSION

Antimicrobial resistance surveillance increasingly relies on homology-based genomic detection of resistance determinants to anticipate therapeutic options. However, novel resistance mechanisms with little or no sequence similarity to known determinants may escape this detection. Gfa173 provides a clear example of this limitation. It encodes a previously undescribed glutathione-dependent fosfomycin-inactivating enzyme with no detectable homology to canonical Fos determinants that would go undetected using sequence-based prediction approaches. While the fitness cost and moderate mobility of Gfa173 might limit its spread, its resistance to PPF inhibition can specifically select this protein among other fosfomycin resistance determinants. Additionally, PPF resistance indicates a catalytic architecture that is fundamentally different from classical FosA and FosB enzymes. Plus, the unexpected restoration of Gfa173 activity in the presence of EDTA supports a clear mechanistic divergence from canonical FosA/B-like enzymes. These proteins require a catalytically essential divalent metal ion (Mn^2+^) that is readily chelated by EDTA, resulting in loss of activity. In contrast, our data indicate that Gfa173 does not depend on a freely exchangeable divalent cofactor for catalysis under the conditions tested although we identified Zn^2+^-cysteine cages. One plausible explanation is that trace divalent metals present in simplified buffer systems interfere with the thiol-dependent catalytic machinery of Gfa173, either by promoting cysteine oxidation or by perturbing the geometry of its cysteine-rich active site^46,47^. Chelation of such trace metals by EDTA may restore a reaction-permissive redox environment^46^. This interpretation is consistent with the behavior of Zn^2+^-cysteine structural complexes, in which zinc is tightly coordinated and does not behave as a labile catalytic metal ion but instead contributes to structural stabilization and proper protein folding^48^. Gfa family members are structurally linked to cysteine-rich, zinc-associated enzymes involved in glutathione-dependent formaldehyde metabolism. However, no formaldehyde-associated resistance phenotype was detected in our assays, in agreement with previous biochemical studies reporting the absence of measurable catalytic acceleration under standard conditions^46,47^. Based on our experimental observations and previous structural and biochemical studies, we therefore propose a putative model for Gfa173-mediated fosfomycin inactivation. Our data indicate that fosfomycin inactivation may occur within a thiol-dependent catalytic mechanism. Despite this mechanistic divergence with metal-dependent FosA-like determinants, the chemical outcome of the reaction may be similar. FosA-mediated inactivation proceeds via nucleophilic attack of glutathione on the epoxide ring of fosfomycin, generating a ring-opened glutathione adduct^49^. In Gfa173, the strict requirement for conserved cysteine residues suggests that epoxide opening may instead be initiated by a protein thiol within the cysteine-rich region, potentially involving the C27–C29–C32 cluster, followed by participation of glutathione. Fosfomycin contains a strained epoxide susceptible to thiol attack, whereas cleavage of the C–P bond is not required for the FosA reaction^49^. Whether Gfa173 ultimately generates the same glutathione–fosfomycin adduct remains to be established experimentally.

The Gfa-like family is widely distributed across Proteobacteria and is predominantly encoded within chromosomal loci, suggesting a role as part of a conserved metabolic route. However, a minority of homologs are embedded within mobile genetic elements, like transposons or integron cassettes. In our dataset, functional analyses revealed that fosfomycin resistance is restricted to a narrow subset of Gfa homologs. Despite conservation of the cysteine-rich architecture and predicted metal-associated fold across all tested proteins, only Gfa173 and two additional homologs identified in distinct *Cellvibrio* species conferred measurable resistance. Structural superposition of representative functional and non-functional homologs confirmed that the overall fold is conserved across the family (Supplementary Figure S2). Thus, conservation of the cysteine clusters alone is insufficient to predict a resistance phenotype. To further investigate this, we selected Gfa^fuscus^, which shares a comparable level of amino acid identity with the functional Gfa^KYGH^ and performed progressive mutagenesis to identify residues capable of restoring resistance. These efforts did not reveal a key determinant, suggesting that resistance cannot be attributed to a single missing catalytic residue. Consistent with this, the evolved Gfa^fuscus^ variant—despite increased sequence identity to functional homologs—did not show improved global structural agreement with Gfa173 in 3D alignments (Figure S3). Altogether, fosfomycin inactivation appears to depend on subtle and globally coordinated structural features that are not readily inferable from sequence alignment alone. This supports the view that functional divergence within the Gfa family reflects distributed structural constraints rather than isolated active-site variation.

Importantly, the existence of numerous Gfa homologs that do not currently confer detectable fosfomycin resistance does not argue against their evolutionary relevance. Rather, these proteins may represent proto-resistance elements within a broader latent resistome^50,51^, retaining a structural framework from which resistance activity could potentially emerge through mutation, altered expression, or mobilization. The failure to restore resistance in Gfa^fuscus^ through a limited set of targeted substitutions suggests that this transition is not governed by a small number of obvious residues and may require more complex evolutionary trajectories.

A limitation of this study is the absence of a complete biochemical characterization of the Gfa173 catalytic mechanism. Although the protein was successfully expressed and purified, we did not identify the reaction product directly, or the kinetic parameters under fully defined reaction conditions. Consequently, the proposed catalytic model is supported by convergent genetic, structural, mutagenesis, docking, and crude-extract inactivation data, but the precise chemical transformation and sequence of catalytic events remain to be established experimentally.

In conclusion, we identify Gfa173 as the founding member of a previously unrecognized family of fosfomycin resistance determinants embedded within mobile integron gene cassettes. Unlike canonical Fos enzymes, Gfa displays a distinct cysteine-dependent, zinc-organized architecture, is refractory to phosphonoformate inhibition. The widespread yet heterogeneous distribution of Gfa-like homologs across Proteobacteria highlights the existence of a latent enzymatic reservoir from which clinically relevant resistance phenotypes can emerge under appropriate selective and genetic contexts. Our findings underscore the limitations of homology-based resistance surveillance and demonstrate that functional screening remains essential to uncover mechanistically divergent resistance determinants. Recognition of Gfa173-like resistance determinants in genomic surveillance will therefore require criteria capable of distinguishing resistance-conferring variants from the broader pool of Gfa-like proteins.

## Data availability statement

All data generated or analysed during this study are included in this article and its Supplementary information. Additional data are available from the corresponding authors upon reasonable request.

## Acknowledgments

We thank Ester Sevillano Gonzalez for her valuable advice and technical support during protein purification and the development of biochemical assays for Gfa173.

## Funding

N.K.: European Union’s Horizon 2020 Marie Skłodowska-Curie grant agreement no. 847635, Instituto de Salud Carlos III, Miguel Servet program CP25/00035

A.P.: Universidad Complutense PhD program

L.O.M.: Ministerio de Ciencia, Innovación y Universidades FPU21/03268 A.H.: Universidad Complutense PhD program

J.A.E.: Ministerio de Ciencia, Innovación y Universidades CNS2022-135857, European Research Council Starting Grant (803375), BBVA Foundation OTR15444, Ministerio de Ciencia e Innovación PID2024-159576NB-I00, EU HARISSA JPI-AMR program PCI2021-122024-2A.

