## Supplemental data for "Gfa, a new integron-borne fosfomycin resistance protein family"

**Table S1. List of bacterial strains used in this study**

|  | Function |
| --- | --- |
| *E. coli* MG1655 | Control strain |
| *E. coli* Dh5α | Control strain, cloning recipient |
| *K. pneumoniae* ATCC23357 | Control strain |
| *P. aeruginosa* PAO1 | Control strain |
| *E. coli* MG1655 pMBA |  |
| *E. coli* MG1655 pMBA-gfa^173^ |  |
| *E. coli* MG1655 pMBA-gfa^KYGH^ |  |
| *E. coli* MG1655 pMBA-gfa^CellV1^ |  |
| *E. coli* MG1655 pMBA-gfa^CellV2^ |  |
| *E. coli* MG1655 pMBA-gfa^PAO1^ |  |
| *E. coli* MG1655 pMBA-gfa^fuscus^ |  |
| *E. coli* MG1655 pMBA-gfa^173^-gfpKO |  |
| *E. coli* MG1655 pMBA-gfa^KYGH^-gfpKO |  |
| *E. coli* MG1655 pMBA-gfa^CellV1^-gfpKO |  |
| *E. coli* MG1655 pMBA-gfa^CellV2^-gfpKO |  |
| *E. coli* MG1655 pMBA-gfa^PAO1^-gfpKO |  |
| *E. coli* MG1655 pMBA-gfa^fuscus^-gfpKO |  |
| *K. pneumoniae* ATCC23357 pMBA |  |
| *K. pneumoniae* ATCC23357 pMBA-gfa^173^ |  |
| *K. pneumoniae* ATCC23357 pMBA-gfa^KYGH^ |  |
| *K. pneumoniae* ATCC23357 pMBA-gfa^CellV1^ |  |
| *K. pneumoniae* ATCC23357 pMBA-gfa^CellV2^ |  |
| *K. pneumoniae* ATCC23357 pMBA-gfa^PAO1^ |  |
| *K. pneumoniae* ATCC23357 pMBA-gfa^fuscus^ |  |
| *P. aeruginosa* PAO1 pBT |  |
| *P. aeruginosa* PAO1 pBT-gfa^173^ |  |
| *P. aeruginosa* PAO1 pBT-gfa^KYGH^ |  |
| *P. aeruginosa* PAO1 pBT-gfa^CellV1^ |  |
| *P. aeruginosa* PAO1 pBT-gfa^CellV2^ |  |
| *P. aeruginosa* PAO1 pBT-gfa^PAO1^ |  |
| *P. aeruginosa* PAO1 pBT-gfa^fuscus^ |  |
| *E. coli* β2163 | Recombination assays |
| *E. coli* β2163 pSWT-attC-gfa^173^-Bottom strand | Recombination assays |
| *E. coli* β2163 pSWT-attC-gfa^173^-Top strand | Recombination assays |
| *E. coli* β2163 pSWT-attC-*aadA7*-Bottom strand | Recombination assays, control strain |
| *E. coli* MG1655 *attB*:*gfp* |  |
| *E. coli* BL21(DE3) | Protein production |
| *E. coli* BL21(DE3) pMAL-closed | Protein production, control strain |
| *E. coli* BL21(DE3) pMAL-gfa173 | Protein production |
| *E. coli* BL21(DE3) pMAL-fosG | Protein production |

**Table S2. List of oligonucleotides used in this study**

| **Name** | **Sequence 5'-3'** | **Function** | **Reference** |
| --- | --- | --- | --- |
| GfafuscusSR-EA-Fw | CAGTGAAgaggcgGTCAGAGATACAAATGCATAATTAGGC | Site directed mutagenesis of gfafuscus | This study |
| GfafuscusSR-EA-Rv | CTCTGACcgcctcTTCACTGCCAGGCCAGCCTTCAAACTG | Site directed mutagenesis of gfafuscus | This study |
| GfafuscusDR-NK-Fw | CTATAGAaacaaaCCCGAAGTTATCGGTGTCCCGCTTGGT | Site directed mutagenesis of gfafuscus | This study |
| GfafuscusDR-NK-Rv | CTTCGGGtttgttTCTATAGGTGCTGATCAAAGGCGAGCC | Site directed mutagenesis of gfafuscus | This study |
| Gfafuscus72F-Fw | CACCAAGtttTTTTGCAAAACGTGTGGCTCGCCTTTG | Site directed mutagenesis of gfafuscus | This study |
| Gfafuscus72F-Rv | TGCAAAAaaaCTTGGTGACGTTTTCTGACGAACCGTA | Site directed mutagenesis of gfafuscus | This study |
| GfafuscusD49N-Fw | TGGTAGTaatCAGTTTCGGTGGTTGTCGGGTGAGCAA | Site directed mutagenesis of gfafuscus | This study |
| GfafuscusD49N-Rv | GAAACTGattACTACCAATGGACGCGCGAGTGCGGAA | Site directed mutagenesis of gfafuscus | This study |
| GfafuscusF25Y-Fw | TCCAATTtacAATTGTCATTGTTCCAAGTGTCGTCGG | Site directed mutagenesis of gfafuscus | This study |
| GfafuscusF25Y-Rv | GACAATTgtaAATTGGACCAAGAGGGCCTTCAATTTC | Site directed mutagenesis of gfafuscus | This study |
| GfafuscusV4M-Fw | GAAGAATGAGCAAGatgACAGGGAAGTGTTTGTGCGAAAC | Site directed mutagenesis of gfafuscus | This study |
| GfafuscusV4M-Rv | AGTTAAGTGATcacTAGTAGGAAGAATGAGCAAGatgACA | Site directed mutagenesis of gfafuscus | This study |
| GfafuscusS48K-Fw | CTCGCGCGTCCATTGGTAaaAATCAGTTTCGGTGGTTGTCGGGTG | Site directed mutagenesis of gfafuscus | This study |
| GfafuscusS48K-Rv | GCATGGCGCGGCTTTCCGCACTCGCGCGTCCATTGGTAaaAATC | Site directed mutagenesis of gfafuscus | This study |
| GfafuscusT113V-Fw | GGCACATATTTTTgttGGTTCAAAGGCAAGCTGGCATCAAATC | Site directed mutagenesis of gfafuscus | This study |
| GfafuscusT113V-Rv | GACATTCAGGGTGGGCCGGAGGCACATATTTTTgttGGTT | Site directed mutagenesis of gfafuscus | This study |
| GfafuscusTR64-66SS-Fw | TCagcagcTACaaaTCGTCAGAAAACGTCACCAAGTTTTTTTGC | Site directed mutagenesis of gfafuscus | This study |
| GfafuscusTR64-66SS-Rv | TGACGATTTGTAgctgctGAGATTTTGCTCACCCGACAAC | Site directed mutagenesis of gfafuscus | This study |
| GfafuscusG64K-Fw | TCagcagcTACaaaTCGTCAGAAAACGTCACCAAGTTTTTTTGC | Site directed mutagenesis of gfafuscus | This study |
| GfafuscusG64K-Rv | TGACGAtttGTAgctgctGAGATTTTGCTCACCCGACAAC | Site directed mutagenesis of gfafuscus | This study |
| pMAL-close-Fw | CCAACAAGGACCATAGATTATGAGGCATCAAATAAAACGAAAGG | Close pMAL vector | This study |
| pMAL-close-Rv | CCTTTCGTTTTATTTGATGCCTCATAATCTATGGTCCTTGTTGG | Close pMAL vector | This study |
| Gfa173-C27A-Fw | GTGagcATTGTAGATTGGACCGAGCGAACCTGTGACTTCG | Site directed mutagenesis of gfa173 | This study |
| Gfa173-C27A-Rv | GTCCAATCTACAATgctCACTGCTCAAAATGCAGGCGCTG | Site directed mutagenesis of gfa173 | This study |
| Gfa173-C32A-Fw | gcgCTGGCATGGTTCAGCTTTTAGAACTCGGGCATCTATC | Site directed mutagenesis of gfa173 | This study |
| Gfa173-C32A-Rv | TAAAAGCTGAACCATGCCAGCGCCTagcTTTTGAGCAGTG | Site directed mutagenesis of gfa173 | This study |
| Gfa173-C8A-Fw | gcTTTGTGCGAAACTGTCGTGTACGAAGTCACAGGTTC | Site directed mutagenesis of gfa173 | This study |
| Gfa173-C8A-Rv | ACGACAGTTTCGCACAAAgcCTTTCCTTTCATTATGTTCA | Site directed mutagenesis of gfa173 | This study |
| Gfa173-C10-Fw | TGTTTGgcCGAAACTGTCGTGTACGAAGTCACAGGTTC | Site directed mutagenesis of gfa173 | This study |
| Gfa173-C10A-Rv | ACGACAGTTTCGgcCAAACACTTTCCTTTCATTATGTTCA | Site directed mutagenesis of gfa173 | This study |
| Int-gcu173Fw | GACGGTATCGATAAGCTTGATATCGTATTCCTCTGGGTGGCATAG | attC cloning into pSWT | This study |
| gBlockR-attC | GCGGCCGCTCTAGAACTAGTGCTGCAGCGTCGACGCCTAA | attC cloning into pSWT | This study |
| gfa-PAO-Fw | GCAGTCGCCCTAAAACAAAGACGCCTTGCAAAGGAGTCAC | Amplify and clone gfaPAO | This study |
| gfa-PAO-Rv | GCTGCAGCGTCGACGCCTAACTCTCTTCTCCTGCTGGAAC | Amplify and clone gfaPAO | This study |
| C74A-Fw | GTTACAAAGTTTTTCgccAACTTGCGGTTCACCTTTAA | Punctual mutation of gfa173 | This study |
| C74A-Rv | TTggcAAAACTTTGTAACATTTTCAGATGATTTATACG | Punctual mutation of gfa173 | This study |
| C77A-Fw | AACCggcTTGCAGAAAAACTTTGTAACATTTTC | Punctual mutation of gfa173 | This study |
| C77A-Rv | TTTCTGCAAAACTgccTCACCTTTAATTAGCACATATG | Punctual mutation of gfa173 | This study |
| pMAL-FosG-Fw | CTCAGCATCATCATCATCATGTGCTCCGAGGATTGAACCA | Clone fosG into pMAL | This study |
| pMAL-FosG-Rv | GTTGTAAAACGACGGCCAGTTTAAAGCCATACTAAGCCCG | Clone fosG into pMAL | This study |
| GFA-to-pMAL-F | CTCAGCATCATCATCATCATATGAACATAATGAAAGGAAAG | Clone gfa into pMAL | This study |
| GFA-to-pMAL-R | GTTGTAAAACGACGGCCAGTTTAAGTATTTGTTTTTCTAAC | Clone gfa into pMAL | This study |
| pMALBB5 | ACTGGCCGTCGTTTTACAACG | Amplify pMAL vector | This study |
| pMALBB3 | ACCATGATGATGATGATGCTGAG | Amplify pMAL vector | This study |
| Int R para backbone | CTTTGTTTTAGGGCGACTGC | Amplify pMBA |  |
| GFP F para backbone | TTAGGCGTCGACGCTGCA | Amplify pMBA |  |
| Gfa173-K116AF | GCTCAgcAGCAACATGGCATAAAATTACTGACTCTG | Site directed mutagenesis of gfa173 | This study |
| Gfa173-S115A-F | GCgCAAAAGCAACATGGCATAAAATTACTGACTCTG | Site directed mutagenesis of gfa173 | This study |
| Gfa173-K116A-R | ATGCCATGTTGCTgcTGAGCCTACGAAAATATGAGCC | Site directed mutagenesis of gfa173 | This study |
| Gfa173-S115A-R | ATGCCATGTTGCTTTTGcGCCTACGAAAATATGAGCC | Site directed mutagenesis of gfa173 | This study |
| Gfa173-S30A-F | TCTACAATTGTCACTGCgCAAAATGCAGGCGCTGGCATGG | Site directed mutagenesis of gfa173 | This study |
| Gfa173-S30A-R | ATTTTGcGCAGTGACAATTGTAGATTGGACCGAGCGAACC | Site directed mutagenesis of gfa173 | This study |
| Gfa173-C29A-F | TCTACAATTGTCACgcaTCAAAATGCAGGCGCTGGCATGG | Site directed mutagenesis of gfa173 | This study |
| Gfa173-C29A-R | ATTTTGAtgcGTGACAATTGTAGATTGGACCGAGCGAACC | Site directed mutagenesis of gfa173 | This study |
| Gfa173-S66A-F | CAgCaGAAAATGTTACAAAGTTTTTCTGCAAAACTTGCGG | Site directed mutagenesis of gfa173 | This study |
| Gfa173-S66A-R | CTTTGTAACATTTTCtGcTGATTTATACGAAGACAAG | Site directed mutagenesis of gfa173 | This study |
| Gfa173-N68A-F | CATCTGAAgcaGTTACAAAGTTTTTCTGCAAAACTTGCGG | Site directed mutagenesis of gfa173 | This study |
| Gfa173-N68A-R | CTTTGTAACtgcTTCAGATGATTTATACGAAGACAAG | Site directed mutagenesis of gfa173 | This study |
| Gfa173-T70A-F | CATCTGAAAATGTTgCAAAGTTTTTCTGCAAAACTTGCGG | Site directed mutagenesis of gfa173 | This study |
| Gfa173-T70A-R | CTTTGcAACATTTTCAGATGATTTATACGAAGACAAG | Site directed mutagenesis of gfa173 | This study |
| Gfa173-R140A | TTGTTTTTGCAACTGCCTCTTCAGACCCTGGCCAATTTTC | Site directed mutagenesis of gfa173 | This study |
| Gfa173-R140A | AGAGGCAGTTGCAAAAACAAATACTTAACAAGGCGTTGCAG | Site directed mutagenesis of gfa173 | This study |
| Gfa173-R43A-Fw | GATGCCGCAGTTCTAAAAGCTGAACCATGCCAGCGCCTGC | Site directed mutagenesis of gfa173 | This study |
| Gfa173-R43A-Rv | AGCTTTTAGAACTGCGGCATCTATCAATAAAAACCAGTTC | Site directed mutagenesis of gfa173 | This study |
| Gfa173-R41A-Fw | CGAGTTGCAAAAGCTGAACCATGCCAGCGCCTGCATTTTG | Site directed mutagenesis of gfa173 | This study |
| Gfa173-R41A-Rv | GGTTCAGCTTTTGCAACTCGGGCATCTATCAATAAAAACC | Site directed mutagenesis of gfa173 | This study |
| Gfa173-K71A-Fw | AAACGCTGTAACATTTTCAGATGATTTATACGAAGACAAG | Site directed mutagenesis of gfa173 | This study |
| Gfa173-K71A-Rv | CTGAAAATGTTACAGCGTTTTTCTGCAAAACTTGCGGTTC | Site directed mutagenesis of gfa173 | This study |
| post-attC-Gfa-Rv | GCGGCCGCTCTAGAACTAGTATCGCTGCAGCGTCGACGCC | attC cloning into pSWT | This study |
| pre-attC-Gfa-Fw | GACGGTATCGATAAGCTTGATATCCAGGCGCTGGCATGGTTCAG | attC cloning into pSWT | This study |
| post-attC-Gfa-Rvbis | GACGGTATCGATAAGCTTGATATCATCGCTGCAGCGTCGACGCC | attC cloning into pSWT | This study |
| pre-attC-Gfa-Fwbis | GCGGCCGCTCTAGAACTAGTCAGGCGCTGGCATGGTTCAG | attC cloning into pSWT | This study |

**Table S3. Antibiotic disks used for antimicrobial susceptibility screening**

| **Antibiotic class** | **Bio-Rad catalog no.** | **Antibiotic** | **Disk content** |
| --- | --- | --- | --- |
| Aminoglycoside | 66148 | Amikacin | 30 µg |
| β-lactam (aminopenicillin) | 68042 | Amoxicillin | 20 µg |
| β-lactam + β-lactamase inhibitor | 66147 | Amoxicillin–clavulanic acid | 20/10 µg |
| Macrolide | 67008 | Azithromycin | 15 µg |
| β-lactam (monobactam) | 66928 | Aztreonam | 30 µg |
| β-lactam (4th-generation cephalosporin) | 66098 | Cefepime | 30 µg |
| β-lactam (3rd-generation cephalosporin) | 66368 | Cefotaxime | 30 µg |
| β-lactam (cephamycin) | 66228 | Cefoxitin | 30 µg |
| β-lactam (1st-generation cephalosporin) | 66218 | Cephalothin | 30 µg |
| Phenicol | 66278 | Chloramphenicol | 30 µg |
| Fluoroquinolone | 68648 | Ciprofloxacin | 5 µg |
| Tetracycline | 66388 | Doxycycline | 30 µg |
| β-lactam (carbapenem) | 67518 | Ertapenem | 10 µg |
| Phosphonic acid antibiotic | 67658 | Fosfomycin | 50 µg |
| Aminoglycoside | 66608 | Gentamicin | 10 µg |
| Aminoglycoside | 66618 | Kanamycin | 30 µg |
| Tetracycline | 66728 | Minocycline | 30 µg |
| Quinolone | 68618 | Nalidixic acid | 30 µg |
| Nitrofuran | 67328 | Nitrofurantoin | 100 µg |
| Rifamycin | 66648 | Rifampin | 5 µg |
| Aminoglycoside | 67418 | Streptomycin | 10 µg |
| Sulfonamide | 68408 | Sulfonamides | 100 µg |
| Tetracycline | 67448 | Tetracycline | 30 µg |
| Glycylcycline | 67398 | Tigecycline | 15 µg |
| Aminoglycoside | 67488 | Tobramycin | 10 µg |
| Folate-pathway inhibitors | 68898 | Trimethoprim–sulfamethoxazole | 25 µg |

**Table S4**. List of the nucleotide sequence of each *gfa* homolog. In blue is the coding sequence.

>*gfa*173

TTAAGTGATCACTAGTAGGAAGAATGAACATAATGAAAGGAAAGTGTTTGTGCGAAACTGTCGTGTACGAAGTCACAGGTTCGCTCGGTCCAATCTACAATTGTCACTGCTCAAAATGCAGGCGCTGGCATGGTTCAGCTTTTAGAACTCGGGCATCTATCAATAAAAACCAGTTCAAATGGCTAAACGGCTCTGAAAACTTGTCTTCGTATAAATCATCTGAAAATGTTACAAAGTTTTTCTGCAAAACTTGCGGTTCACCTTTAATTAGCACATATGAAAACAAACCTGAAATTTTAGGTATTCCTCTGGGTGGCATAGAAGGCATTACTGAAGCCAATATTGAGGCTCATATTTTCGTAGGCTCAAAAGCAACATGGCATAAAATTACTGACTCTGTGCCACAATATGAAAATTGGCCAGGGTCTGAAGAGGCAGTTAGAAAAACAAATACTTAACAAGGCGTTGCAGCACCGTTCCGGCTTCGCCTCCACTGGACAGTTTGTAAGTTGCGCTTTTGTGGTTTTGCTTCGCAAAAGTATTCCACAACGCGCAACTTACAAACTGCCGCTGAACTTAGCG

>*gfa*^KYGH^

ATGATTGGAAAATGTTTGTGTGAAGCGGTTGAGTACACGATCGATGGTGCCTTAGGGCCGGTTTACAACTGCCATTGTTCAAAGTGTCGCCGTTGGCATGGGGCTGCATTTAGAACGCGTGCTTCAATTAATAAAAATCAATTCACTTGGATTAAAGGTGAGGCGAATTTATCCAGCTATAAATCGTCTGATTATGTCACTAAATTTTTTTGTAAAACCTGTGGTTCTCCCCTTATCAGTACTTATGACAATAAGCCCGATATTCTAGGTGTTCCACTCGGTGGTATTGAGGGGATTGAGCACGTGGAAGTGCAAGCACATATTTTTGTGGGTTCAAAAGCGAGTTGGTATGAGATTGCTGATTTAGCGCCTCAGTACGACGCCTGGCCAGGGAGTGAAGAGGCTGTCCGTGAAACTAATCAATAG

>*gfa*^CellV1^

ATGAGTTTAATGAAAGGAAAATGTTTGTGCGAGTCGGTTCAGTATGAAATTACCGGTTCGCTTGGTCCAATTTATAATTGCCATTGCTCAAAGTGCAGAAGGTGGCATGGTTCAGCATTTAGAACTCGTGCATCCATAGATAAAAATCAATTCAGGTGGATTAGTGGTTCGGAAAATTTGTCTTCTTACAAATCATCTGAAAATGTTACAAAGTTTTTCTGTAAAACCTGTGGCTCACCACTCATTAGCACTTATGAAAATAAACCTGACGTTTTGGGTATTCCGCTGGGTGGCATTGAAGGAATCCCTGAAGTTAATGTAGAAGCAAATATATTTGTTGGTTCAAAAGCATCGTGGTTCAAAATTACAGACTCAGTGCCACAATTTGAAGCTTGGCCAGGTTCAGAGGAAGCAGTTAGAAAAACGAACAATTCAAACTCTGAGTGA

>*gfa*^CellV2^

ATGAAAGGCAGTTGTCTTTGTGGTGGTATTAAGTTTAGCTTCAAAAAATTTGACTCTAAAATCGCTAATTGCCACTGTTCTATGTGCAGAAAATTTAGCGGTTCGGCATACGGAACTTTCGGACCTGTAAATTCTGAAAACATGAATTGGGAATCTGGGTTTGAGCTAATCAAAATTTTCAACTCTTCTCAAAACGCTCATCGAGGCTTTTGTTCAATTTGTGGTTCAAGTATTTTTTATCGCTTAACAAGCAAAAATTCTCCATTTGAAATTGCACTTGGGTTATTGGAAGCCGAGCCAACCGAATCAGTTTCAGCTAACATATTTTGTGCGTATCGCCCAAATTGGCCACGGAACGTAGATGAGATTCCAAATTTTGAAGAATCAAGAACGGTTTAA

>*gfa*^PAO1^

TTGCAAAGGAGTCACGAAATGGATCGATTCACCGGCGGCTGCCTCTGCGGCAAGGTCCGCCTGGTGGCGAGCGGCCGCCCCTACCGGGTCGGGCTCTGCCACTGCCTCGACTGCCGCAAGCACCACGGCGCGCTATTCCACGCTTCGGCGATCTTTCCCGAGGAGGCGGTGAGCATAGAAGGCGAGACCCGCGACTATGCCGGGCGTTTCTTCTGTCCGCAGTGCGGCTCCTCGGTATTCTCGCGCAGCGCCGACGAGATCGAGGTCAGCCTCGGCGCCCTCGACGCTCCCGACCGGTTCCAGCCCACCTACGAACTCTGGACCGTCCGCCGCGAGGGCTGGTTGCCGGCGTTTCCGCTGGCACGGCACTATGAAAGGGATCGCGAAGGCGACGGACGCAGCGAAGAGTGA

>*gfa*^fuscus^

ATGAGCAAGGTAACAGGGAAGTGTTTGTGCGAAACAGTGAGCTATGAAATTGAAGGCCCTCTTGGTCCAATTTTCAATTGTCATTGTTCCAAGTGTCGTCGGTGGCATGGCGCGGCTTTCCGCACTCGCGCGTCCATTGGTAGTGATCAGTTTCGGTGGTTGTCGGGTGAGCAAAATCTCACACGGTACGGTTCGTCAGAAAACGTCACCAAGTATTTTTGCAAAACGTGTGGCTCGCCTTTGATCAGCACCTATAGAGACCGTCCCGAAGTTATCGGTGTCCCGCTTGGTGGTCTGGAAGGTGACATTCAGGGTGGGCCGGAGGCACATATTTTTACTGGTTCAAAGGCAAGCTGGCATCAAATCACAGACTCACTGCCACAGTTTGAAGGCTGGCCT GGCAGTGAATCGAGGGTCAGAGATACAAATGCATAA

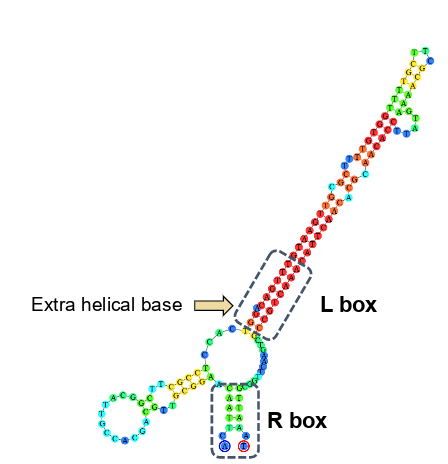

**Supplementary Figure S1. Representation of the *gfa173* *attC* site**. The *gfa* gene is associated with a well-structured *attC* site featuring a canonical L box, R box, and extrahelical base (arrow), supporting its mobilization via integron recombination. Free energy:ΔG = -26.37 kcal/mol, indicating a highly stable secondary structure compatible with efficient integrase recognition.

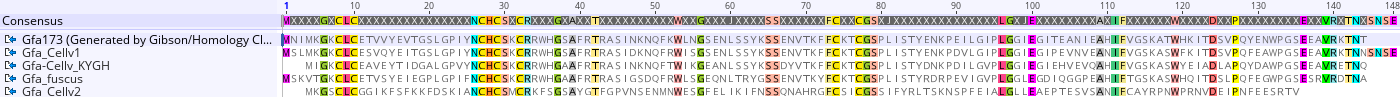

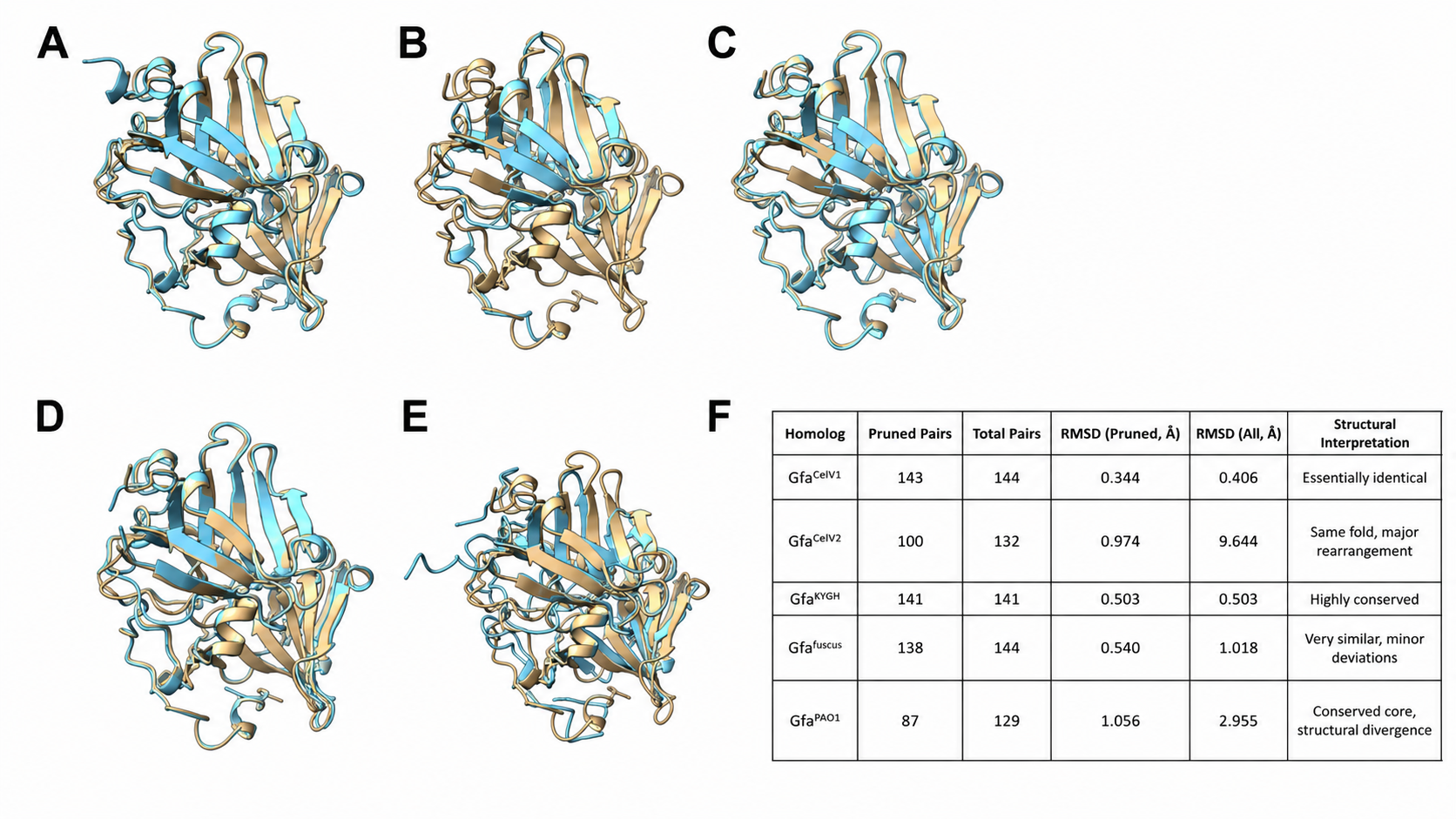

**Supplementary Figure S2. Alignment and structural superposition of Gfa homolog dimers.**

**Top.** Amino acid alignment of all Gfa homologs. **Bottom.**(A–E) Ribbon representations of Gfa173 (reference dimer, blue) structurally aligned with selected homolog dimers (beige) using the MatchMaker algorithm in ChimeraX. (A) Gfa^CellV1^, (B) Gfa^CellV2^, (C) Gfa^KYGH^, (D) Gfa^fuscus^, and (E) Gfa^PAO1^. In all cases, full dimer assemblies were superimposed. Despite functional divergence, all homologs retain the characteristic β-sandwich architecture and overall dimeric organization of the Gfa family. Functional homologs (Gfa^CellV1^ and Gfa^KYGH^) display near-identical backbone alignment with the reference, whereas non-functional or divergent homologs exhibit increasing peripheral deviations while preserving the conserved core fold. (F) Summary of dimer structural alignments, including number of aligned residue pairs and RMSD values (pruned and total). Backbone RMSD values confirm high structural conservation at the dimer level despite differences in fosfomycin resistance phenotype.

**Supplementary Figure S3. Sequence and structural comparison of Gfa173 with Gfa^fuscus^ and evolved Gfa^fuscus^.
(A)** Amino-acid sequence alignment of the original Gfa^fuscus^ protein and the evolved Gfa^fuscus^ variant, highlighting the limited number of substitutions introduced during stepwise mutagenesis. **(B)** Pairwise amino-acid identity matrix comparing Gfa173, Gfa^fuscus^, and evolved Gfa^fuscus^. Values indicate percentage amino-acid identity. **(C)** Structural comparison of Gfa173 with the original and evolved Gfa^fuscus^proteins. Structural superpositions are shown for Gfa173 versus Gfa^fuscus^ (left) and Gfa173 versus evolved Gfa^fuscus^ (right). The table summarizes the structural alignment metrics, including the number of pruned and total aligned residue pairs and the RMSD values calculated for pruned and full alignments. Together, these analyses indicate that the evolved Gfa^fuscus^ variant remains highly similar to Gfa173 after pruning, while showing greater overall structural divergence than the original Gfa^fuscus^ homolog.

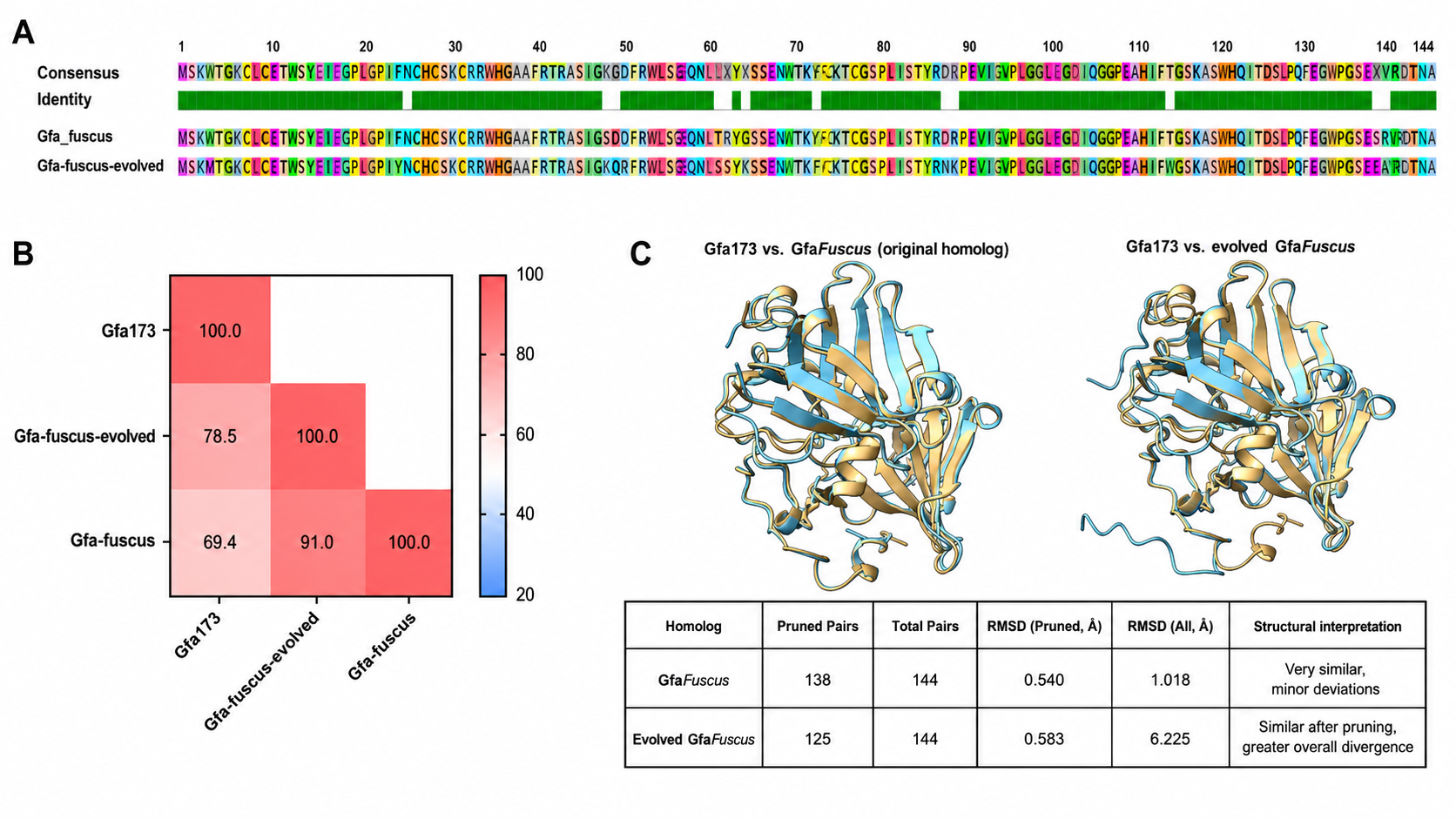

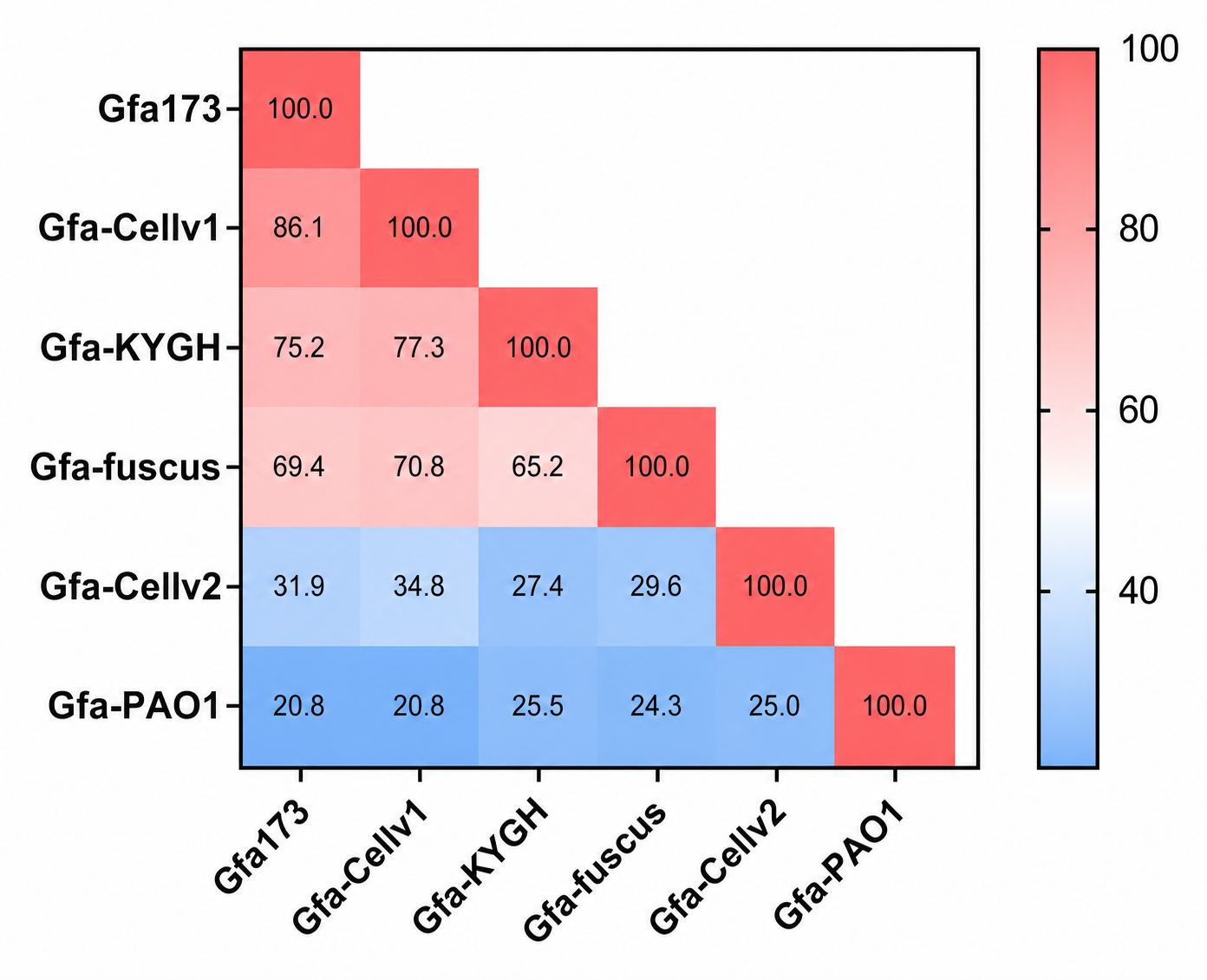

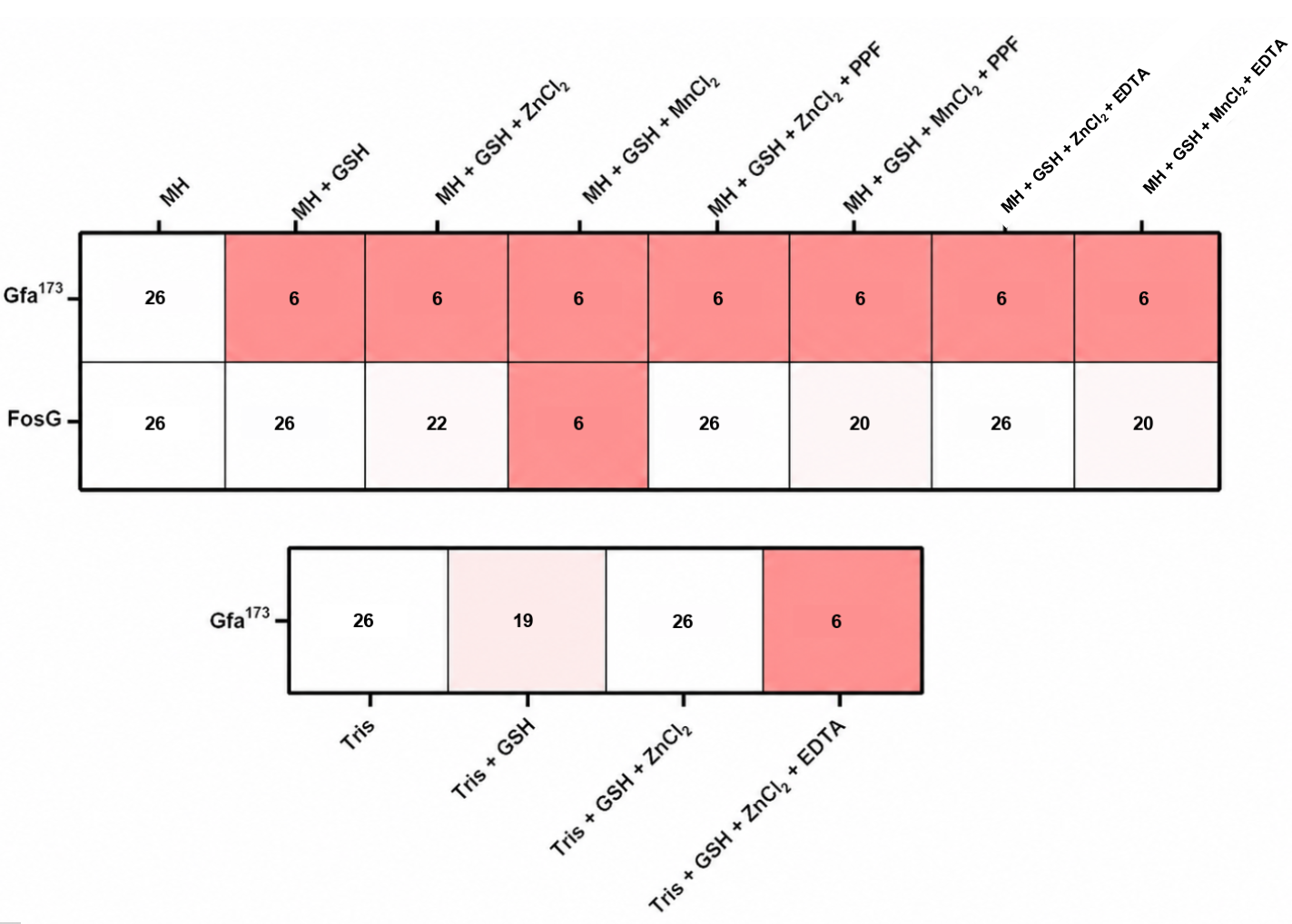
